# Cryo-EM-based structural insights into supramolecular assemblies of γ-Hemolysin from *Staphylococcus aureus* reveal the pore formation mechanism

**DOI:** 10.1101/2022.09.14.507916

**Authors:** Suman Mishra, Anupam Roy, Somnath Dutta

## Abstract

γ-hemolysin (γ-HL) is a hemolytic and leukotoxic bicomponent β-pore-forming toxin (β-PFT), a potent virulence factor from *Staphylococcus aureus* Newman strain. In this study, we performed single particle cryo-EM of γ-HL in a lipid environment. We observed clustering and square lattice packing of octameric HlgAB pores upon membrane bilayer, and an octahedral superassembly of octameric pore complexes, that we resolved at resolution 3.5 Å. Our atomic model further demonstrated the key residues involved in hydrophobic zipping between the rim domains of adjacent octameric pore complexes, thus providing first evidence of additional structural stability in PFTs upon membrane lysis. We also observed lipid densities at the octahedral and octameric interfaces, providing critical insights into the lipid-binding residues involved for both HlgA and HlgB components. Furthermore, the hitherto elusive N-terminal region of HlgA has also been resolved in our cryo-EM map and an overall mechanism of pore formation for bicomponent β-PFTs is proposed.

## INTRODUCTION

*Staphylococcus aureus* (*S. aureus*) is a gram-positive bacterium typically associated with nosocomial infections. Depending on the site of infection, *S. aureus* can cause a variety of complications, a few of which include meningitis, endocarditis, brain abscesses, skin and soft tissue infection, bacteraemia, osteomyelitis, and pneumonia. *S. aureus* has gradually emerged as a life-threatening pathogen since the discovery of its resistance to multiple generations of antibiotics, e.g., penicillin, vancomycin, tetracycline, and methicillin, thereby significantly increasing morbidity and cost burden^1–5^. This “superbug” primarily seizes the entire host defense mechanism through its appalling arsenal of virulence factors which is not limited to but essentially comprises immune hijacking and modulatory toxins, superantigens and pore-forming toxins (PFTs)^5,6^. PFTs are dimorphic proteins-they are first secreted as watersoluble monomeric components, which then oligomerize and form amphipathic pores on host cell membranes with the hydrophobic patches exposed to the bilayer acyl chains and the hydrophilic regions lining the periphery^7^. Most of the virulent strains of *S. aureus* including community-associated methicillin-resistant *S. aureus* (CA-MRSA) and hospital-acquired methicillin-resistant *S. aureus* (HA-MRSA), express six cytolytic toxins that are haemolytic and/or leukotoxic^5^. Among the six, only α-hemolysin (Hla)^8^, β-hemolysin^9^ and δ-hemolysin^10^ are composed of a single polypeptide^11^. The other three cytolysins, namely γ-Hemolysin (γ-HL), leukocidin (Luk), and Panton-Valentine leukocidin (PVL), are bicomponent β-PFTs^11,12^. Based on sequence similarity with the two Luk components, monomeric toxin components for all such bi-component β-PFTs are either classified as a class F or class S component^12^. Bi-component β-PFTs require synergistic oligomerization of a class F component and a class S component, i.e., γ-HL, Luk, and PVL are composed of LukF (HlgB) and Hlg2 (HlgA/HlgC), LukF and LukS, and LukF-PV and LukS-PV as class F and class S component proteins, respectively^11^. These heterodimers associate with each other to form a ring-shaped pre-pore structure which then develops a β-barrel, forming a pore^13–15^.

Several structural and biochemical studies on oligomeric γ-HL and its individual Hlg2, HlgC, and LukF components have previously elaborated, (i) the most stable oligomeric pore complex^13,16,17^, (ii) the relative stoichiometries of individual monomeric components in the stable oligomeric pore complex^13,14^, (iii) the significance of γ-HL in destruction of erythrocytes and leukocytes during *S. aureus* infection^18^, (iv) its recently discovered link to TRPV1-associated surge in physical pain^19^ and, (v) DARC-mediated necrosis^20,21^. Interestingly, stable oligomeric pore complexes of mono-component α-hemolysin and bicomponent γ-HL were reported to be induced in the presence of 20 percent 2-Methyl-2,4-pentanediol (MPD), a widely used crystallizing agent^13,22^. MPD binding site in the rim domain of γ-HL crystal structure correlated with the lipid head group (di-propanoyl phosphatidyl choline, or DiC_3_PC) binding sites previously predicted for the monomeric LukF component, however, only MPD and not DiC_3_PC was able to induce oligomerization^13,23–25^. However, solution state structure of stable oligomeric pore complexes of γ-HL in the context of a lipid membrane remains uncharacterized. There is limited understanding regarding the crucial steps of PFT assembly, such as conversion of pre-stem to stem, the formation of transmembrane β-barrel via stem domains, and the oligomerization of cap domains^7,26,27^. Importantly, the fate of such pore complexes upon lysis of lipid membranes has not been studied thus far.

Several recent cryo-EM based structural studies have attempted to illuminate the mechanism of oligomeric assembly and pore formation by PFTs in a near-native environment. In addition to mapping the atomic structures of most stable oligomeric states, solution state studies so far also have had the advantage of trapping intermediate pre-pore assemblies^28,29^, capturing the pore complex in a model membrane^29–32^, as well as unearthing rather uncommon oligomeric stoichiometries^33,34^. Therefore, we chose to characterize the solution state structure of bicomponent PFT, γ-HL in a physiologically relevant membrane microenvironment by employing cryo-EM single particle analysis. For this structural study, we targeted HlgAB pore complexes, comprising HlgA (LukS/Hlg2) and HlgB (LukF) components from *S. aureus* (strain Newman), owing to its robust virulence phenotype. We purified the two toxin components, HlgA and HlgB, tested the functional activity, and obtained transmembrane pore complexes induced by large unilamellar liposomes constituting phosphatidylcholine and cholesterol for NS-TEM and cryo-EM-based experiments. Interestingly. we were able to visualize, for the first time for wildtype PFTs, a higher order octahedral supramolecular assembly of octameric HlgAB pore complexes stabilized by hydrophobic zipping of rim domains, that we resolved to global resolution of 3.5 Å. Our atomic model does not only enunciate the structural basis of this novel phenomenon, but it also throws light on the atomic structure of the hitherto uncharacterized N-terminal region and its possible supporting role in pre-stem to stem conversion of the adjacent component during pore assembly. Finally, through our structural study, we also provide detailed experimental evidence for several crucial lipid-binding residues for both HlgA and HlgB components that might contribute to the initial tethering of monomers to a lipid surface.

## RESULTS

### Preliminary characterization of recombinant bicomponent HlgAB pore complexes from *Staphylococcus aureus* Newman strain

In this study, we aimed to characterize the 3D structure of bicomponent HlgAB pore complexes in the presence of lipid environment at cryogenic conditions using single particle cryo-EM. To achieve our target, we initially performed overexpression and two-step protein purification of the two individual components, HlgA (LukS component) and HlgB (LukF component). SEC elution profile indicated HlgA and HlgB eluted around 18-20 ml, and SDS PAGE showed both the components were highly homogeneous (Figure 1A-B)The identity of both the proteins was also confirmed by peptide mass fingerprinting of trypsin-digested components(Figure S1). Previously, it was reported that α-HL and γ-HL could oligomerize into a stable pore complex upon treatment with 15-20% of 2-Methyl-2,4-pentanediol (MPD), a common crystallizing agent^13,22^. Therefore, we incubated equimolar concentrations of the two components-HlgA, and HlgB in 20% MPD (v/v) at 37 °C to observe the formation of any stable oligomer. The purity and stability of MPD-treated HlgAB was visualized by SDS-PAGE, where a prominent band above the standard molecular weight of 250 kDa was observed, that might correspond to the estimated molecular weight of ~280 kDa for octameric HlgAB (Figure 1C). The stable oligomeric state for several PFTs, including the bicomponent octameric state of HlgAB, is known to be resistant to SDS and could be observed via SDS-PAGE^13^. Room temperature NS-TEM raw micrographs of the MPD incubated species as well as their reference-free 2D class averages illustrated the formation of uniform, homogeneous oligomeric pore complexes with a wide central cavity (Figure 1C). However, in our current study, we were interested to observe the oligomeric states of HlgAB in the presence of a real lipid environment. Therefore, we selected large unilamellar vesicles (LUVs) constituting equimolar concentrations of egg-PC and cholesterol as a model membrane for our study.

**Figure 1.**
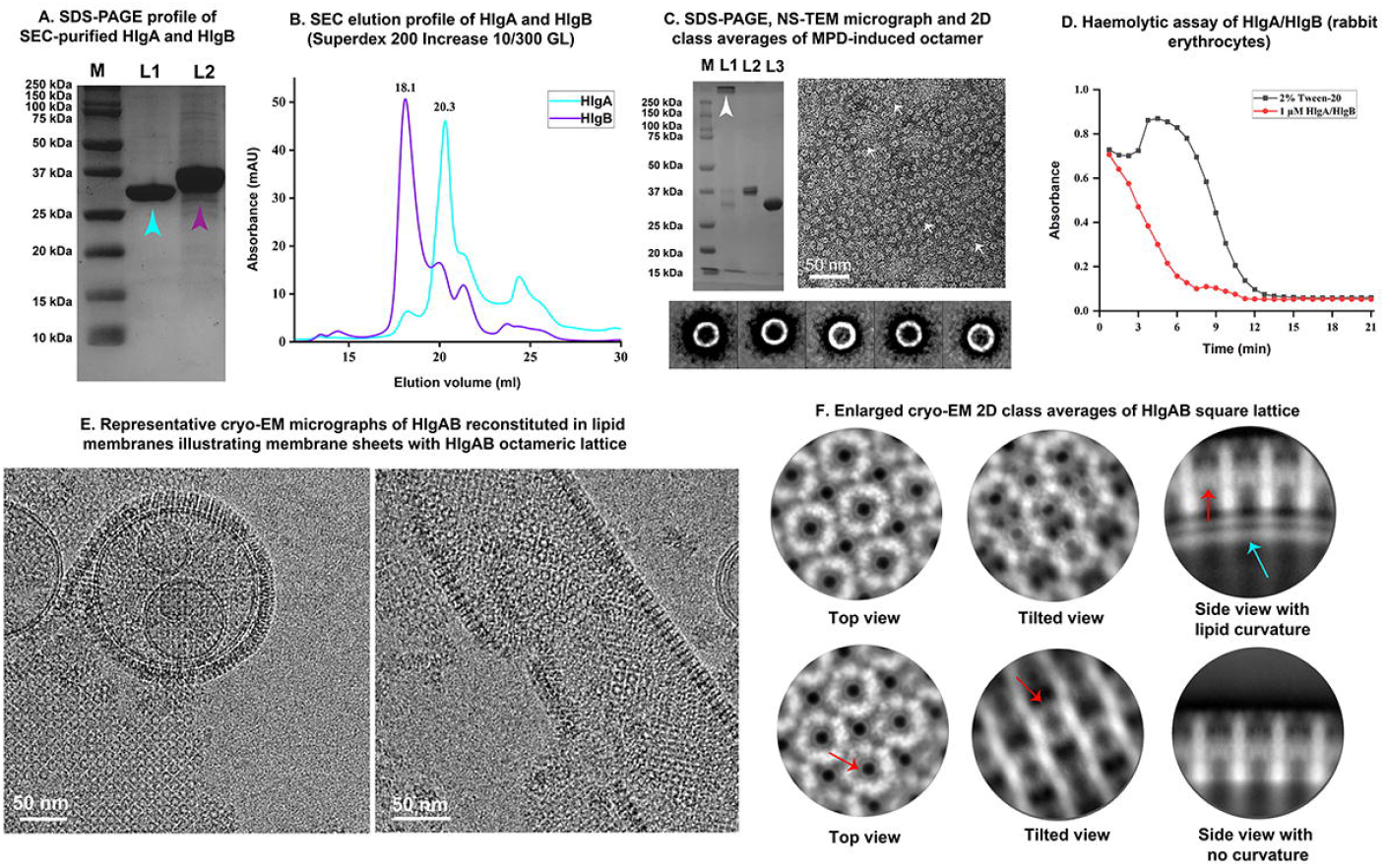
Purification, functional characterization, and cryo-EM analysis of bicomponent γ-HL. **A.** 12% SDS-PAGE analysis of purification of recombinant HlgA (L1) & HlgB (L2). Lane M; Protein size standards (Bio-Rad Precision Plus Protein™ Dual Color Standards). Lanes 1 and 2 correspond to gel filtrated fractions of HlgA (fraction volume at 20.3 ml) and of HlgB (fraction volume at 18.1 ml), respectively. **B.** Gel filtration chromatography analysis of HlgA (cyan) and HlgB (purple) on Superdex 200 increase 10/300 GL column. SEC elution profiles of 32kDa 6xHis-tagged HlgA & 34 kDa 6xHis-tagged HlgB in terms of *A*_280_ are shown in solid lines. **C**. 10% SDS-PAGE analysis upon treating equimolar concentrations of freshly purified recombinant HlgA and HlgB with 20% (v/v) MPD. A prominent SDS-stable oligomeric band (L1; white arrowhead) is observed above the standard molecular weight of 250 kDa. L2 and L3 represent HlgB and HlgA monomeric bands. Negatively stained-transmission electron microscopy (NS-TEM) micrograph of HlgA and HlgB treated with 20% MPD illustrate the formation of homogeneous oligomeric pore complexes (white arrows). Corresponding 2D class averages indicate ring-like pore complexes. **D.** Haemolytic assay of rabbit erythrocytes was performed in experimental triplicates for functional characterization of the two protein components. 2% Tween-20 detergent was used as positive control. Absorbance was measured at 620 nm. Black and red lines indicate curves of 2% Tween-20 and HlgA/HlgB treated erythrocytes, respectively. **E.** Representative cryo-EM micrographs of HlgAB reconstituted in lipid membranes illustrating square lattice packing of octameric pore complexes upon globular LUVs (left) and ruptured membrane sheets (right). **F.** Particles corresponding to lattice arrays were subjected to 2D classification with an enlarged box size. Selected cryo-EM class averages illustrate different views of lattice packing as well as octamer. Cyan arrow indicates liposome curvature, red arrows indicate different orientations of transmembrane β-barrel.

Again, upon incubation of the two protein components HlgA and HlgB with LUVs and observing under NS-TEM, we could observe formation of protein complexes and disruption of the membrane surface that was distinct from our control liposomes (Figure S2A-B, S3). In accordance with a previous study on HlgAB^17^, we could also observe the tendency of these protein complexes to form larger clusters upon ruptured membranes (Figure S2B, S3). Using rabbit erythrocytes, we further confirmed that the two purified proteins also possessed hemolytic activity (Figure 1D) that we attribute to the formation of HlgAB pores decorating the erythrocyte surface as observed under NS-TEM (Figure S2C, S4).

### Self-association tendency of bicomponent HlgAB octameric pore complexes and its square lattice arrangement upon LUVs

Interestingly, upon egg-PC-cholesterol LUVs treated with the two PFT components, we encountered several remarkable features of this protein complex. After visualizing membranes fully studded with HlgAB pore complexes resembling a lattice-like arrangement under NS-TEM (Figure S2B, S5), we acquired cryo-EM datasets to investigate HlgAB pore complexes in context of lipid membranes in cryogenic conditions. We further discerned that HlgAB pore complexes tended to self-associate (Figure S6), forming local clusters, larger non-specific aggregates, or two-dimensional lattice upon lipid membranes. Notably, we could observe such crystalline lattice arrangement on intact spherical liposomes as well as ruptured arcs and sheets of lipid membranes (Figure 1E, S6). This tendency of HlgAB made three-dimensional reconstruction of individual pore complexes extremely challenging with limited side views and crowding of particles. However, we performed reference-free 2D classifications and observed several novel insights from the resultant 2D class averages (Figure 1F). First, we could identify individual pore complexes in the crystalline sheets to be octameric assemblies packed predominantly in a square lattice (p4m) array. We did not observe any deviation from square lattice in raw micrographs as well as 2D class averages (Figure 1E-F, S6, S7B). Second, we were able to obtain distinct class averages for tilted views and side views of 2D lattice array (Figure 1F) which correlate with previous cryo-EM studies on protein lattice arrays^35^. Different orientations of the β-barrel domain were clearly distinguishable in all 2D classes (Figure 1E), also suggesting that matured transmembrane pore complexes participated in the lattice formation. Third, we were able to observe the phenomenon of curvature in protein lattice array at a molecular level. Raw micrographs as well as 2D class averages revealed that HlgAB pore complexes forming lattice upon intact LUVs tended to acquire their respective curvature (Figure 1E-F, S8). Such intricate artificial protein-lipid associations have been reported in several past studies in which a 2D array of proteins was formed upon planar lipid sheets^36,37^, however, in our study, the square lattice arrangement was also remarkably observed to be maintained upon liposome curvature. Upon reconstructing an electron density map from a single 2D class average corresponding to top views of square lattice, we were able to grossly fit the atomic model (PDBID: 3B07) and compare the biological assembly of octameric pore complexes of HlgAB to the crystallographic assembly as reported for LukF-Hlg2 hetero-octamer (PDBID: 3B07) (Figure S7). As opposed to the antiparallel arrangement of pore complexes within a plane in the case of crystallographic symmetry (Figure S7A), we could observe that in physiological state, HlgAB pore complexes tended to lie parallel within a plane as they adopted square lattice packing (Figure S7B). Clustering of HlgAB pore complexes and square lattice formation suggested that the matured oligomeric pore complexes were partially destabilized upon rupturing of membranes. This phenomenon of association of pore complexes upon rupturing lipid membranes caught our attention, and we further targeted larger self-association of HlgAB pores to unravel the fate of such PFTs upon membrane lysis.

### Room temperature NS-TEM and single particle analysis cryo-EM reveal octahedral supramolecular assembly of octameric HlgAB pore complexes

Under NS-TEM, we were able to visualize some small clustering of protein-lipid complexes which might have been multiple oligomeric HlgAB complexes clustered around ruptured membranes. To our surprise, NS-TEM raw micrographs of LUVs-treated HlgA and HlgB not only contained these octameric pore complexes but also several orientations of a hitherto unidentified species that was ~20 nm in diameter (Figure S8A-B). Upon performing reference-free 2D classifications of small protein lipid clusters, we were able to distinguish primarily three distinct orientations of this species, namely, “corolla” view that appeared to be a hexamer in architecture, “lantern” view composed of two pore complexes equatorially extending from a ~20 nm long meridian, and “cross” view representing four units perpendicular to each other and one pore complex extending outside the plane (Figure S8B). A time-dependent study of HlgA and HlgB in lipid environment revealed that such species were being formed within 30 minutes post incubation along with the routine octameric species, suggesting a higher assembly of pore complexes (Figure S9). To investigate whether these species were only associated with membrane-based study, we also attempted to observe this phenomenon upon treatment of HlgA and HlgB with 20 percent of MPD. Remarkably, we were able to observe homogeneously distributed particles, predominantly in the “corolla”, and “lantern” orientations under NS-TEM (Figure S10A). The resultant reference-free 2D classification revealed similar features as described for the species observed in lipid environment (Figure S10B). However, the crystal structure of octameric HlgAB was determined previously in the presence of MPD, but the study did not observe the presence of such super assemblies^13^. This is the first time we have showed the phenomenon of super assembly of HlgAB in the presence of both MPD and liposomes. The homogeneity in particle size and presence of distinct orientations of this species encouraged us to further target these particles for a three-dimensional reconstruction using single particle analysis.

However, related single particle cryo-EM studies of HlgAB were performed in presence of a lipid environment to also identify the role of lipids in pore formation.

To enunciate the molecular architecture of this species in native state, we manually picked the particles corresponding to the abovementioned distinct orientations (Figure 2A, Figure S11A). Our cryo-EM 2D class averages, also consistent with NS-TEM data, revealed highly resolved structural features through which we were able to interpret that this species was a supramolecular assembly of HlgAB- and, actually comprised matured octameric HlgAB complexes with their transmembrane β-barrel domain facing inwards (Figure 2B, Figure S11B). We calculated an *ab initio* model without imposition of any symmetry with these particles (Figure S12) and determined the 3D structure of HlgAB at 3.5 Å resolution (gold standard FSC, 0.143 cut-off) that was fit for subsequent atomic model building (Figure 2C, S12, S13B). Furthermore, local resolution estimation using Blocres predicted most regions of the map corresponding to a resolution of ~3.2 Å (Figure 3A). Interestingly, we observed that this supramolecular assembly was an octahedron with four octameric bicomponent HlgAB pore complexes lying in a plane perpendicular to each other and two pore complexes extending meridianally at 180° (Figure S13A). Furthermore, the atomic model explained the molecular aspects of the formation of HlgAB octahedral supramolecular assemblies, and to further elaborate upon the solution state structure of bicomponent HlgAB pore complexes. Densities corresponding to atomic model’s two individual components HlgA and HlgB with their respective side chain fitting also illustrated high resolution features throughout our super assembly electron density map (Figure 3B-C).

**Figure 2.**
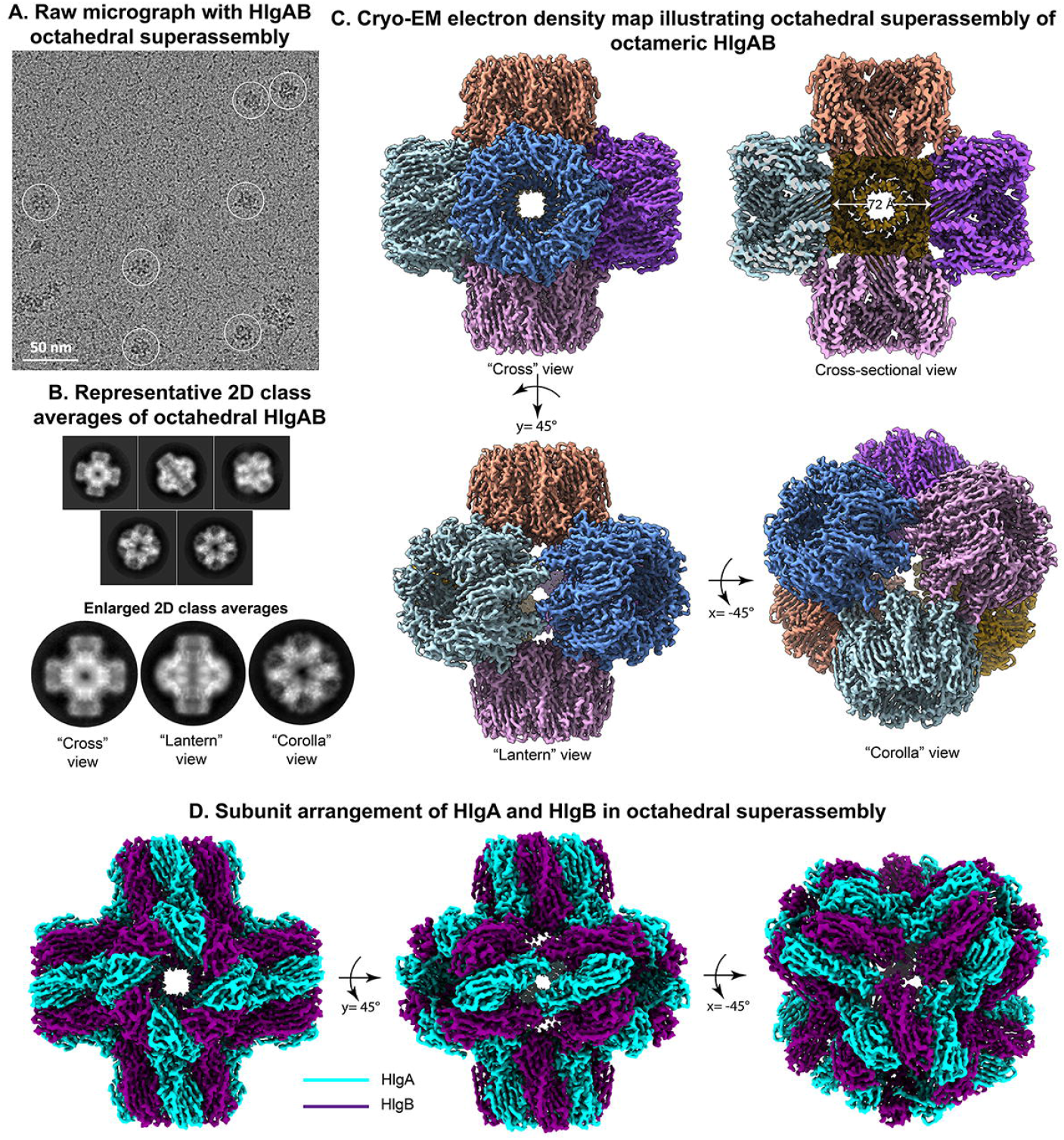
Structural characterization of HlgAB octahedral supramolecular assembly of octameric pore complexes using cryo-EM. **A.** Raw micrograph with HlgAB octahedral superassembly particles encircled in white circle. **B.** Representative 2D class averages of octahedral superassembly of HlgAB (top) and enlarged 2D classes illustrating cross view (four octamers perpendicular to each other in a plane), lantern view (two octamers at 180° and two octamers extending from the centre), and corolla view (hexamer in appearance). Bottom panel shows 2D class averages of negatively stained octahedral superassembly of HlgAB. **C.** Cryo-EM electron density map illustrating the octahedral superassembly of octameric HlgAB at a global resolution of 3.55 Å. Six different octameric HlgAB pore complexes are coloured differently for visualization of supramolecular assembly. Cross, lantern, and corolla views are related by rotation along two axes. Cross sectional view (top right) demonstrates internal rim-to-rim arrangement of the superassembly. **D.** Subunit arrangement of the two components, HlgA (cyan) and HlgB (purple) within octahedral superassembly architecture. Each octamer consists of alternate arrangement of four units of HlgA and four units of HlgB protomers. A total of 48 toxin protomers contribute to the octahedral superassembly.

**Figure 3.**
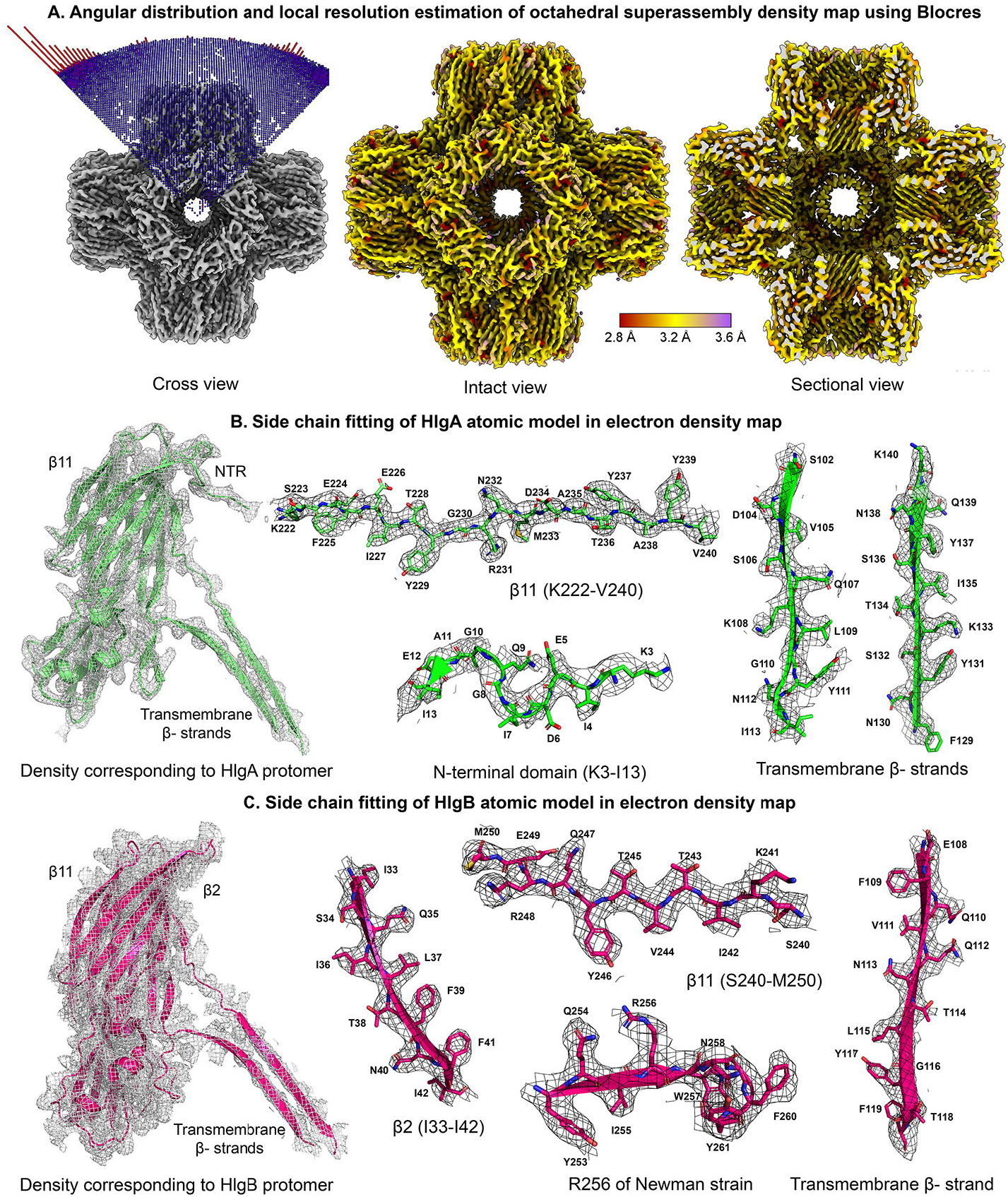
Local resolution estimation and side chain fitting of atomic models of bicomponent HlgAB in octahedral superassembly electron density map. **A.** Angular distribution plot (left) illustrates uniform angular sampling in the particle stack contributing to the final reconstruction. Local resolution estimation using Blocres (centre and right). Colour scale for local resolution is between 2.8-3.6 Å. Frontal view (centre) and sectional view (right) of the octahedral density map indicate the local resolution of major densities to be ~3.2 Å. The β-barrel is also resolved to a resolution of 3.2 Å. **B.** Electron density corresponding to the HlgA component (green) is shown in mesh view (extreme left). Side chain fitting for β11(K222-V240), previously unmodeled N-terminal domain of HlgA (K3-I13), and two transmembrane β-strands (S102-I113, and F129-K140) has been presented. **C.** Density corresponding to the HlgB component (pink) is shown in mesh view (extreme left). Side chain fitting for β2(I33-I42), β11(S240-M250), a β-loop-β region containing R256 residue unique to Newman strain (Y253-F260), and a transmembrane β-strand (E108-T118) has been presented.

### Structural and molecular chemistry of inter octameric pore complex interaction in octahedral super-assembly

Herein, we provided the cryo-EM structure of the octameric pore of bicomponent γ-HL embedded inside its octahedral super assembly complex, which is the first atomic-resolution cryo-EM structure of a higher order supramolecular self-assembled state of any wild-type pore forming toxins (PFTs) in the presence of lipid environment reported till date (Figure 2, Video S1). Such typical octahedral supramolecular hierarchy had a unique topological similarity with common inorganic complexes, such as SF_6_. In comparison to such complexes, the six ligand sites were occupied with six identical octameric pore HlgAB complexes (Figure 2C, S14A-B). From imaginary central point of octahedral topology, four HlgAB pore complexes lied on the four sides of octahedral square plane whereas the other two HlgAB octamers were perpendicular to the square plane and opposite to each other in orientation (Figure 2C, S13A). The cap segments of each pore complex were directed outwards whereas the transmembrane pore composed of sixteen anti-parallel β-strand of stem segments (two β-strand from four stem segments of HlgA and HlgB of octameric HlgAB complexes respectively) directed towards the imaginary central point in octahedral complex (Figure 2C, S13A, S14C). These led to the structural appearance of a hollow cubic nano-cage inside octahedral super assembly having an inner volume of ~373 nm^3^ (Figure 2C).

A detailed investigation of octahedral super-assembly of bi-component γ-HL (HlgA, and HlgB) predicted that mushroom-shaped octameric pore complex attached via nearby four other octameric pore complexes in any square plane present perpendicular to that reference octamer in octahedral topology (Figure 4A). Each octameric pore complex perpendicular to the square plane shared contact surfaces with a total of four neighboring octameric pore complexes that lied in the square plane via the rim domains (Figure 4A, S13A). In such a geometry, we observed that each γ-HL octameric pore complex shared two stable hydrophobic contact with each of the four HlgAB octamer present in a square plane, perpendicular to that of reference octamer. Thereby each HlgAB octameric pore complex shared a total of eight identical contact zones with four other pore complexes (Figure 4A). These inter-octameric rim domain contacts were quite unique since each contact face was symmetrically organized in such a way that one subunit of bi-component γ-HL of any octameric pore complex favored strong hydrophobic clustering of aromatic and steric-zipper residue interactions between Y68, P185, Y190, V240 of HlgA and Y72, A202, Y203 of HlgB vertically (180°) with another component of γ-HL alternatively (Figure 4B-C, S14C-D). Thereby, the entire octahedral superassembly was stabilized by a total of 24 interfacial rim domain contacts between HlgA_poreX_ and HlgB_poreY_ (where octameric pore complex X lies perpendicular to pore Y). Interestingly, the superassembly interface was enriched with the most favorable hydrophobic zipper forming Tyr residues (Y68, Y72, Y190, Y203, Y239) and aided by a few significant sticker-zipper dry hydrophobic clusters formation between Y190 (HlgA)-Y203 (HlgB), P185 (HlgA)-Y72 (HlgB), V240 (HlgA)-Y203 (HlgB) & Y68 (HlgA)-A202 (HlgB) (Figure 4B-D, S14D).

**Figure 4.**
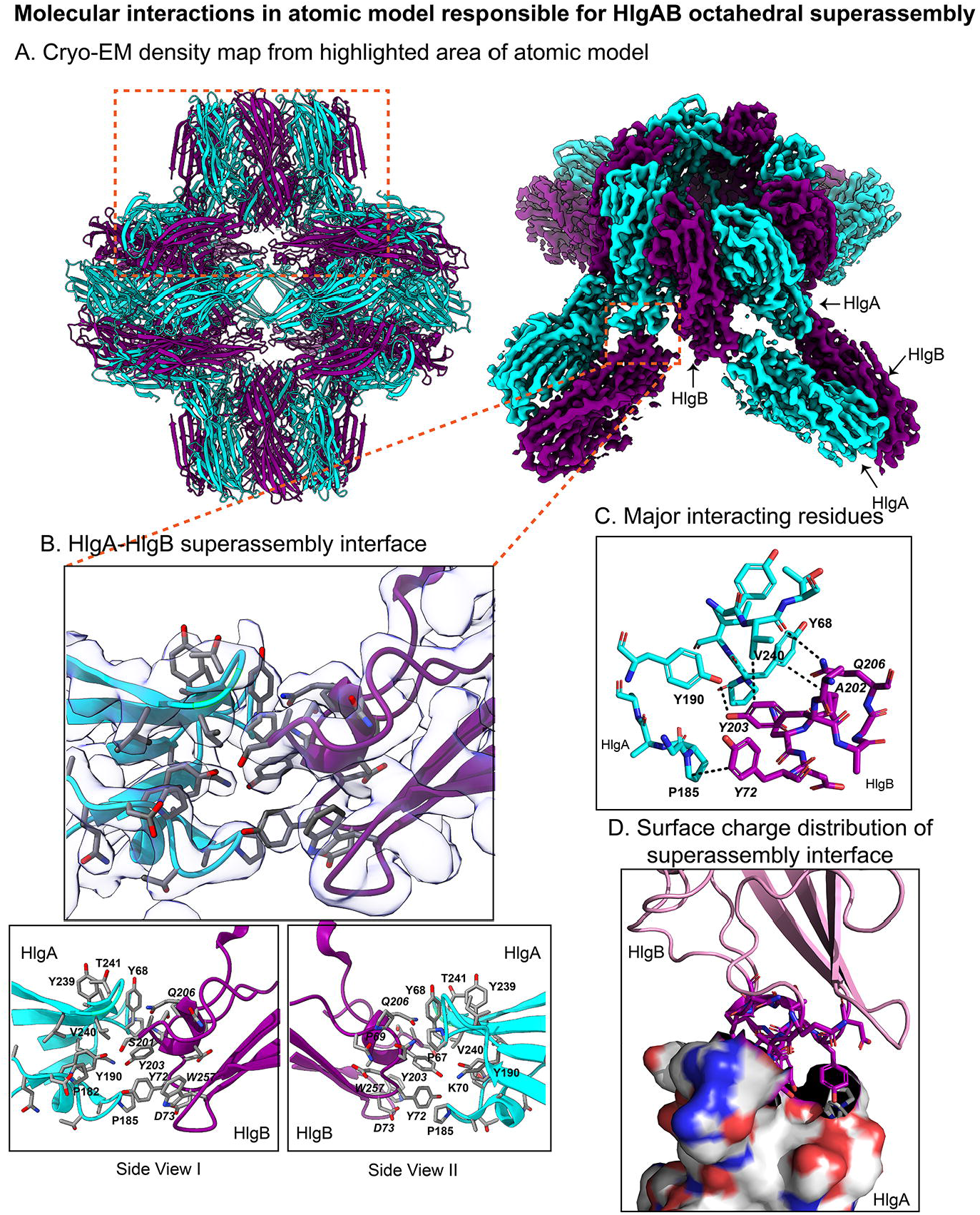
Protein-protein interaction between γ-hemolysin components HlgA and HlgB in the octahedral supramolecular assembly. **A.** The cartoon representation of atomic model (left) of the octahedral super-assembly of γ-hemolysin HlgAB octamers where the constituent individual protomeric subunits are represented in different colours (HlgA in cyan and HlgB in violet). The stereo-representation of the cryo-EM electron density map of selected region of atomic model (right) further predicted the subunit arrangements (HlgA and HlgB) with one octameric HlgAB pore complex inside octahedral topology of bi-components γ-hemolysin HlgAB super-cluster. Each octameric HlgAB pore complex shares two alternative subunits for the interfacial stabilization with adjacent octamers and the rim domains of HlgA (total 24 subunits) get internally neutralized by rim domains of HlgB (total 24 subunits) located at adjacent octamers. **B.** The inter-octameric subunits interactions responsible for octahedral super-assembly formation with highlighted critical amino acid residues (stick model) fitted within translucent electron density map located at the bottom segments of rim domains for both protomers (HlgA/HlgB from highlighted octamer with HlgB/HlgA from adjacent octamers present perpendicular to the reference octamer). Two different views (side views-I and II) illustrate the stereo representation of bottom segment of such rim domains (HlgA-HlgB super-assembly interface) containing aromatic hydrophobic amino acids Y68, Y72, Y190, Y203, Y239 and W257 and other hydrophobic non-aromatic residues P67, P89 and V240 and polar amino acids K70, D73, S201, Q206 and T241, respectively. **C.** The ball and stick model representation of residues participating in interfacial inter-octameric subunits interactions between Y190(HlgA)-Y203(HlgB), P185(HlgA)-Y72 (HlgB), V240 (HlgA)-Y203 (HlgB), Y68 (HlgA)-A202(HlgB), and V240 (HlgA)-Q206 (HlgB). **D.** The surface charge distribution of bottom segment of rim domain of HlgA with the bottom segment of rim domain of HlgB represented in cartoon model.

### Investigating the critical lipid/cholesterol binding segments inside the octahedral HlgAB super-cluster

From our investigation, we could not find self-assembled states where cap domain of one pore complex interacts with another pore complex^28^. Instead, we observed self-assembled array of octameric pore complex on bilayer support and large assemblies of octameric pore complex apart from octahedral superassembly. How did the octameric pore complexes remain stable in water after lysis of lipid membrane where the hydrophobic residues located at the rim domains would face additional level of solvent-exposed instability, remained an enigma. Therefore, the finding of lipid density over/inside the octahedral super-assembly could provide several new insights into identifying the prior lipid-binding sites. This could also predict the critical amino acids responsible for the early event of binding onto lipid surface, as they must possess high affinity of lipid/cholesterol. We observed fuzzy density of lipid/cholesterol through each inter-connected rim segments (rim HlgA_poreX_ vs. rim HlgB_poreY_ and rim HlgA_poreX_ vs. rim HlgA_poreY_) from individual octameric pore complex symmetrically close to super-assembly interface formed by hydrophobic sticker-zipper interactions^38–40^ (Figure 5A, S16B, S17). The amino acids Y68, Y237, Y239, T241, R242, R244, D248 belonging to rim domain of HlgA were proximal to such extra fuzzy density and formed 24 protein-lipid interfaces of HlgA-HlgA (rim HlgA_poreX_ vs. rim HlgA_poreY_) very close to superassembly interface (Figure 5A, S16B, S17). In a similar topological fashion, 24 HlgA-HlgB lipid interfaces (rim HlgA_poreX_ vs. rim HlgA_poreY_) were predominated by the residues D181, P185, L245 of HlgA and Y72, D73, W257 of HlgB with extra fuzzy density of lipid/cholesterol (Figure 5A, S16B, S17). The eight vertices (interface of HlgBporeX/ HlgB_poreY_/ HlgB_poreZ_) could also suffer from a partial topological frustration inside the closed octahedral geometry. We observed a partial lipid/cholesterol density located near residues N258, F260 and Y261 from three adjacent HlgB respectively at the eight vertices (Figure 5A, S16B, S17). This could provide additional protection to the top part of barrel consisting of hydrophobic residues from solvent exposure. Interestingly, the lipid-interacting residues observed at the superassembly interface strongly correlated with the theoretical predictions by Assignment aNd VIsualization of the Lipid bilayer (ANVIL) algorithm to be proximal to the assigned membrane bilayer for crystal structure of LukF-Hlg2 hetero-octamer (Figure S15)^41,42^. A detailed analysis of protein-lipid interfaces inside each octameric pore complex has been further depicted in the subsequent section.

**Figure 5.**
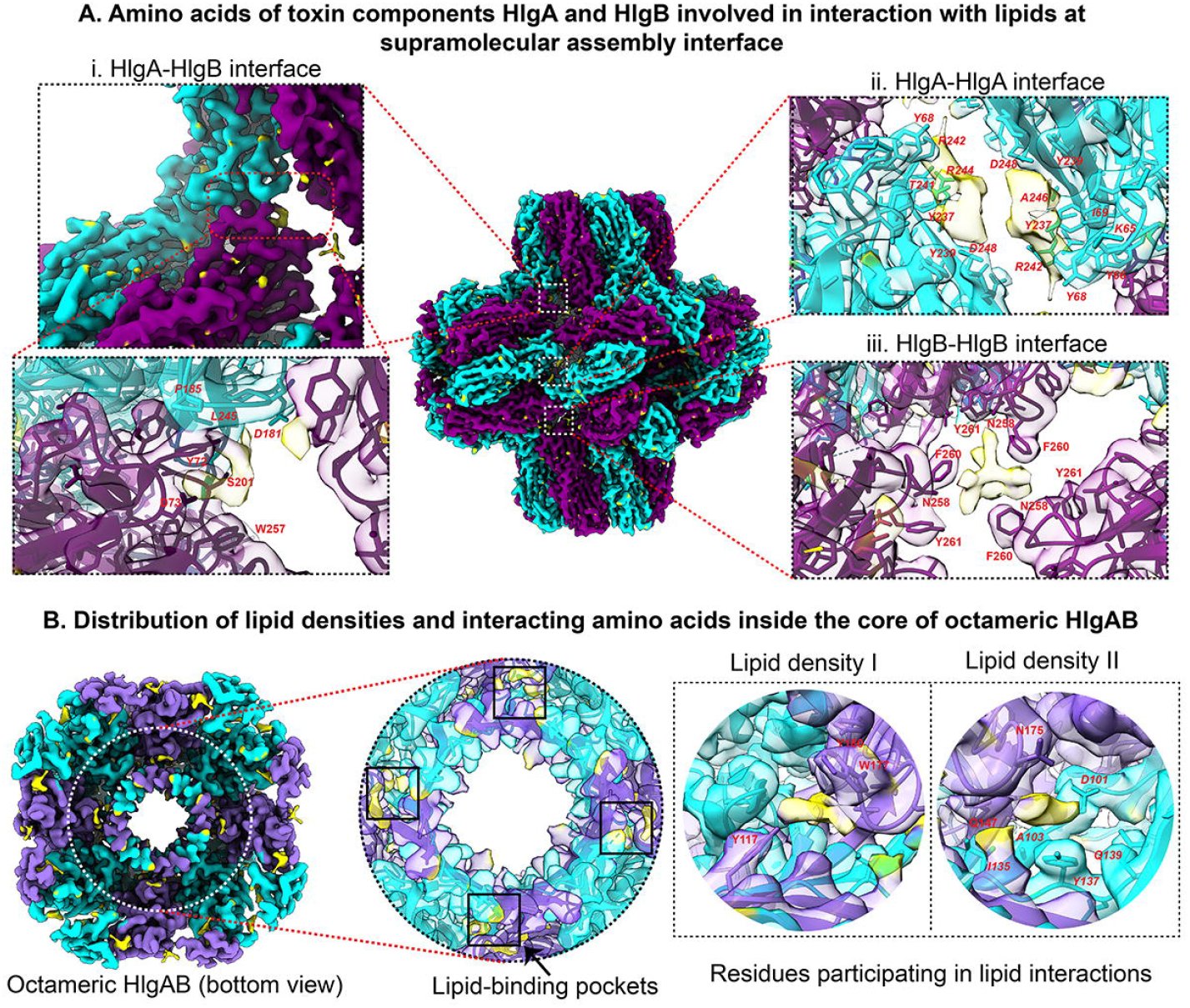
Lipid-interacting residues of HlgA and HlgB components at supramolecular assembly interface and within the core of an individual octameric pore complex of γ-HL. **A.** Amino acids of bicomponent HlgAB involved in interaction with lipids at the superassembly interface. Three different sites of extra densities possibly corresponding to lipids were observed at these interfaces. These sites are marked with white dotted rectangles and enlarged to visualize the neighbouring residues. Extra densities are coloured in yellow, HlgA model and corresponding map densities are coloured in cyan, HlgB model and corresponding map densities are coloured in purple. Residues in italics belong to HlgA, and those in bold belong to HlgB. (i) HlgA-HlgB interface: D181, L245, P185 (HlgA); S201, Y72, D73, W257 (HlgB). (ii) HlgA-HlgA interface: K65, Y66, Y68, I69, Y237, Y239, T241, R242, R244, D248 (HlgA). (iii) HlgB-HlgB interface: N258, F260, Y261 (HlgB). **B.** Distribution of lipid densities and interacting residues inside the core of octameric HlgAB is shown. Electron density map corresponding to one individual bicomponent octameric pore complex in octahedral assembly has been color coded as mentioned above. Lipid-binding cleft has been encircled in white and enlarged to visualize the interacting residues. Two extra densities were observed. The first lipid density has Y117, W177, Y180 (HlgB) in vicinity. The second lipid density has D101, A103, I135, Y137, Q139 (HlgA); Q147, N175 (HlgB) in vicinity.

### Novel insights into the atomic model of octameric bicomponent HlgAB pore complex in octahedral super assembly

A detailed understanding of cryo-EM structure of γ-HL pore complex has further revealed that the bicomponent γ-HL self-assembled into hetero-octameric structure as conformationally most stable geometry. Whereas the octahedral super-assembly state may be highlighted as a native geometrical state in near physiological condition after rupturing of lipid membranes by octameric pore complexes. The structural anatomy of octameric pore complex was composed of alternate arrangement (pseudo-C8) of four subunits each of HlgA and HlgB. The height and outer diameter of octameric pore complex were 71.5 Å (Cα’s of HlgA/G120 and HlgB/K16) and 103.7 Å (Cα’s of C-terminal HlgA/K280 to HlgA/K280) respectively. Whereas the inner pore diameter was measured to be 42 Å (Cα’s of N-terminal HlgB/K16 to HlgB/K16) (Figure S16A). The overall distribution of secondary structural elements in the protomeric HlgA and HlgB inside octameric pore complex shared three distinct structural domains (cap, rim, and trans-membrane β-barrel stem domain) (Figure S16B). Both tertiary structural topology and secondary structure-based multiple sequence alignment suggested that protomeric HlgA and HlgB evolved a similar secondary and tertiary structural geometry (Figure S18). The overall topological features of protomeric HlgA and HlgB resembled ellipsoid geometry having an extended arm containing the stem domain responsible for transmembrane β-barrel formation (Figure S18A).

Furthermore, we superimposed the solution state cryo-EM structure of bicomponent γ-HL octameric HlgAB pore complex (*S. aureus* Newman stain) with the MPD-induced crystal structure of γ-HL octamer of LukF & Hlg2 (RMSD= 0.74 Å for 15200 Cα atoms) (Figure S19). This clearly suggested that even in near physiological conditions, the octameric pore complex was the preferred structural symmetry. Furthermore, the protomeric HlgA and HlgB components from our structure also possessed a large conformational rigidity upon comparison with monomeric HlgA (PDBID: 2QK7), and HlgB (PDBID: 1LKF) (RMSD= 0.704 Å for overlapped 1417 Cα atoms for HlgA, and RMSD= 0.796 Å for overlapped 1670 Cα atoms for HlgB) (Figure S19). We did not observe any significant conformational shift of the segments located at the bottom of rim domains (RMSD=0.532 for HlgA, 0.458 for HlgB) in a structural comparison with γ-HL hetero-octamer of LukF and Hlg2 (PDBID: 3B07) (Figure S19). The overall stability came from the hiding of solvent-exposed hydrophobic residues located in lipid binding rim domain for the two components of γ-HL without having any large dynamic change in the overall structural topology of octameric HlgAB. This further suggested that the formation of octahedral super-assembly might be obvious/natural both in near physiological condition as well as native microenvironments.

Even at atomic resolution of 3.5 Å, the transmembrane β-barrel stem density was partially missing between residues I113-F129 for HlgA, F119-T137 for HlgB in electron density map upon comparison with γ-HL octamer crystal structure (LukF & Hlg2), where the height and diameter of observable β-barrel stem domain are 30 Å (Cα’s of HlgA/K140 and HlgA/S128) and 25.5 Å, respectively (Figure S15D, S16A, S20A). The top and bottom segments of transmembrane β-barrel enriched with hydrophobic residues were further stabilized by intra- and inter-stem β-hairpin interactions (ion pairs and hydrophobic interactions) among adjacent protomers (Figure S20B-C). Furthermore, a strong quaternary complex (hydrophobic belt) was formed by aromatic amino acids (Y111, I113, Y131, F129 of HlgA and Y117, F119, T137, F139 of HlgB) at the termini of our observable stem domain adjacent to the glycine and serine-rich stem segments evolved for both the protomers (Figure S20D). From our EM map, we did not observe any fuzzy/strong lipid/cholesterol density surrounding the hydrophobic surfaces of top part of β-barrel (Figure 5B). The formation of specific supramolecular assembly (octameric pore complexes in octahedral topology) might further protect top segment of transmembrane β-barrel composed of a belt of hydrophobic residues from polar solvents. A large abundance of small and flexible glycine and serine residues present in the bottom part of transmembrane β-barrel composed of four β-hairpin stem segments of HlgA and HlgB might contribute to the observed dynamic nature of β-barrel segments (Figure S18B).

In an observation similar to other *S. aureus* PFTs including homo-heptameric α-hemolysin^8^ (PDBID: 7AHL) and bi-component octameric LukGH^43^ (PDBID: 4TW1), the electron density of N-terminal amino latch density of one subunit (HlgB) of bicomponent HlgAB was missing in cryo-EM density map of HlgAB octamer whereas N-terminal of another component (HlgA) spanned symmetrically throughout the pore (Figure S21). This could suggest that one N-terminal amino latch of bicomponent PFTs was mostly disordered and remained exposed to bulk water, whereas the other four N-terminal segments from another component (HlgA) projected toward nearby HlgB subunit maintaining rotational symmetry (Figure S21A). We observed that the amino acid, E5 of each N-terminal segment of HlgA formed a salt-bridge interaction with N40 residue present in β2 segment of the cap domain of adjacent HlgB (Figure 6A-B).

**Figure 6.**
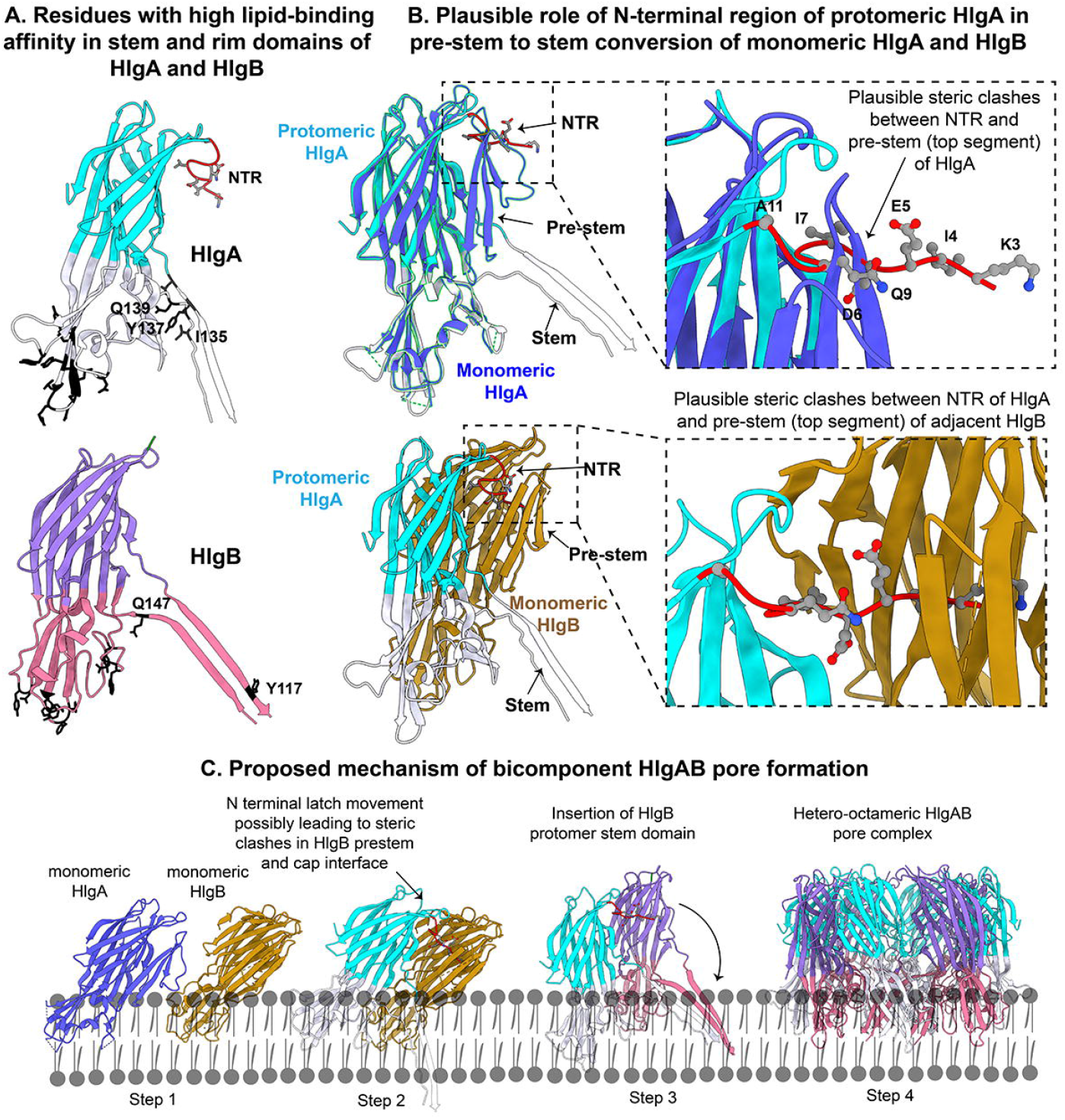
Insights into critical lipid-binding residues in the rim and stem domains and plausible role of N-terminal region of HlgA in pre-stem to stem conversion for both HlgA and HlgB monomers. **A.** Critical amino acids responsible for early-stage binding on lipid surface and the positioning of lipid-binding residues in stem domains of both the protomers. Residues present on stem domain might influence pre-stem to stem conversion upon lipid binding. **B.** Pre-stem unfolding could also be modulated by steric clashes between N-terminal region of HlgA and pre-stem domains of both HlgA and HlgB monomers (PDBID of monomeric HlgA-2QK7, and monomeric HlgB-1LKF). **C.** Proposed mechanism of bicomponent HlgAB pore formation. Step 1-Inclined monomeric HlgA and HlgB embed upon the lipid environment. Rim as well as pre-stem domains simultaneously get exposed to lipid surface. Step 2-N-terminal latch movement from HlgA possibly leads to steric clashes between HlgA as well as HlgB pre-stem and cap interfaces, this could be one of the reasons of pre-stem to stem conversion that leads to insertion of protomer stem domain of HlgB. Step 3-Membrane insertion of stem domain. Steps 2 and 3 could be concerted. Step 4-Subsequent oligomerization leads to formation of a functional hetero-octameric pore complex.

Importantly, we also observed additional fuzzy densities within the HlgAB octameric pore architecture. The residues D101, A103, I135, Y137, Q139 in vicinity of these fuzzy densities belonged to the stem domain of HlgA. Whereas these fuzzy densities (lipid/cholesterol) were located at both large, disordered globule of rim domain (residues: N175, W177 and Y180) as well as stem domain (residues: Y117 and Q147) for HlgB (Figure 5B), which might also correspond to predicted lipid/cholesterol binding sites (Figure S15, S16B).

## DISCUSSIONS

Since the advent of twentieth century, PFTs have garnered a lot of scientific attention owing to their active involvement in bacterial virulence and pathogenesis. Deciphering the diverse pore-formation mechanisms for different families of PFTs still remains the cornerstone of this research. In the backdrop of multidrug resistance, γ-HL from *S. aureus* acts as a great model for structural studies targeting bicomponent β-PFTs for the following two reasons: (1) bicomponent PFTs, such as Leukocidin and Panton Valentine Leukocidin share a significant sequence and structural similarity with γ-HL^12^, and (2) γ-HL has the highest hemolytic and leukotoxic activities among all cytolytic toxins of *S. aureus* and contributes significantly to tissue necrosis during *S. aureus* infection^18,20,21,44^.

In retrospect, our cryo-EM based study throws light on several novel aspects of γ-HL machinery. A key highlight from the study is the novel observation of the supramolecular assemblies of PFTs post membrane lysis, accounted by hydrophobic zipping between the otherwise membrane-embedded rim domains. To the best of our knowledge, this is the first such report in any class of PFTs-thus illuminating the fate of these dimorphic PFTs after rupturing the target lipid membranes (Figure S22). Coalescence of independent γ-HL octameric pore complexes resulting in an octahedral supramolecular assembly is another fascinating example of hydrophobic zipping stabilizing larger assemblies, previously seen for the concentric β-barrel pores of aerolysin^28^ and lysenin^31^. However, for such concentric β-barrel pores, dimerization due to hydrophobic and uncharged residues preceded post-prepore and quasi-pore steps of pore formation^28,31^. In contrast for γ-HL, the octahedral superassembly of octameric pore complexes stabilizing rim domains outside lipid environment was observed to be a product of pore formation and membrane lysis. Consequentially, we hereby anticipate more bacterial PFTs to adopt such protein folding after host membrane lysis.

Recently, several high resolution cryo-EM-based structural studies have targeted PFTs in detergent^28,32,45^, lipid nanodiscs^46,47^, and liposome environment^29,48–50^ to gain structural insights into key residues proximal to the lipid environment. However, this is the first such study on a bicomponent β-PFT in which we provided structural insights into the critical lipid-binding residues. At a global resolution of 3.5 Å, extra densities observed for both the toxin components in our electron density map could correspond to phosphatidylcholine/cholesterol. This does not only strongly correlate with previously reported MPD-binding sites^13^, but also experimentally validates the computational predictions for the membrane spanning regions of γ-HL using ANVIL algorithm^41^. It was interesting to note that γ-HL pore complexes possessed several strong lipid densities even after membrane lysis, suggesting that the residues discovered at such lipid-binding pockets might synergistically play an important role in membrane insertion. From our current investigation, we also acknowledged the dynamic nature of the bottom part of transmembrane β-barrel which was disordered in solution state after lysis of lipid membranes. Glycine and serine-rich bottom part of lytic pore domain could get additional stabilization inside the dry hydrophobic lipid core of the membrane, suggesting that complete transmembrane domain of octameric pore complex of HlgAB might be resolved only upon adherence to lipid bilayer.

Furthermore, our high-resolution density map further empowered us to model the previously unstructured N-terminal domain of HlgA. We found the atomic structure of this N-terminal domain extremely intriguing, since the finding of N-terminal segment symmetrically spanning along the octameric pore complex correlates with the previously reported crystal structures for α-HL and the bicomponent β-PFT LukGH. This evidence suggests a common oligomerization pathway and pore formation mechanism for these bicomponent PFTs of *S. aureus* which are also known to form mixed oligomers, which will serve ground for our future studies on bicomponent β-PFTs.

Finally, to decode the plausible role of lipid-binding residues found to be located in the stem domain, we compared both the monomers with their respective protomeric states. Among the abovementioned residues located at stem domain of HlgA, we identified Y137, Q139 (located in the bottom of pre-stem) and I135 (middle of pre-stem) contributed significant interactions with the hydrophobic surfaces of cap domain. Similar hydrophobic contacts with the cap domain were observed for the residues Q147 (bottom part of pre-stem) and Y117 (top part of pre-stem) in the stem domain of HlgB (Figure S23, S24). Hence, binding of those residues with lipid surface might provide an instability factor for pre-stem with respect to cap domain, promoting early stage unfolding and conversion of pre-stem to stem. This could only be possible if the individual monomeric states bind to lipid surface partially horizontal to its crystallographic axis. Such relative orientational preference of monomeric components was further supported by a recent molecular dynamics study^51^. In addition, the superimposition of monomers upon protomers further elucidated that the critical positioning of N-terminal segments from adjacent protomeric HlgA might influence in destabilizing the surfaces of top segments of cap and pre-stem by a strong steric interference on the nearby hydrophobic and polar residues present in the top part of both monomeric subunits (HlgA and HlgB) respectively (Figure 6B, S23, S24). Thus, binding of pre-stem domain on lipid surface (in association with binding of rim domain to lipid bilayer) and a very critical conformational reorganization of N-terminal segments might possess a structural cooperative effect on pre-stem to stem conversion which was one of the prime steps in pore formation associated with many β-PFTs^7^ (Figure 6C).

In summary, through this study, we have elaborated upon the phenomenon of supramolecular assembly of pore-forming toxins after membrane lysis and its underlying mechanism. The critical lipid binding residues inside the rim domains of HlgA and HlgB components and the modeling of elusive N-terminal region of HlgA also serve as key takeaways that might further illuminate the overall mechanism of pore formation and geometrical consequence of such lethal oligomeric pore complexes.

## METHODS

### Cloning, overexpression, and two-step protein purification of bicomponent HlgAB

Both HlgA (NCBI-ProteinID: BAF68590, UniProt ID: A0A0H3KAA5) and HlgB (NCBI-ProteinID: BAF68592, UniProt ID: A0A0H3KAB8) genes without their respective signal peptides were codon-optimized for expression in *E. coli* BL21 (DE3) and cloned into pET-26b(+) vector (Genscript, USA) with a C-terminal 6X His-tag incorporated for immobilized metal affinity chromatography.

Transformation of both the recombinant plasmids was individually carried out into *E. coli* BL21 (DE3) strain cells to further test the overexpression of the two desired proteins. The transformed cells (resistant to kanamycin) for HlgA and HlgB were then separately grown in Luria Broth (HiMedia Laboratories, LLC) at 37°C to an optical density (at 600 nm (OD600)) of 0.6-0.8. Recombinant proteins (HlgA and HlgB) overexpression was induced by 0.5 mM Isopropyl-b-D-thiogalactoside (IPTG) at 25°C for 16 hours. Cultures were then centrifuged at 4000 rpm for 10 min.

This pellet was then freshly resuspended in lysis buffer (buffer composition-20 mM Tris-HCl buffer pH 7.8, 150 mM NaCl), followed by probe sonication for 30 minutes. The cell lysate post sonication was then centrifuged at 10000 rpm for 40 minutes.

The supernatant solution after this step was then allowed to bind overnight to Ni–nitrilotriacetic acid (NTA) agarose beads pre-packed column, pre-equilibrated with equilibration buffer (buffer composition-20 mM Tris-HCl buffer pH 7.8, 150 mM NaCl). The column was further washed with a wash buffer (buffer composition-20mM Tris-HCl buffer pH 7.8 and 150 mM NaCl) containing gradient imidazole concentration between 20 mM-100 mM, followed by elution of the protein with 300 mM imidazole concentration.

The eluted proteins from Ni-NTA column were then buffer-exchanged and concentrated to 5 mg/ml using Amicon^®^ Ultra centrifugal filter with cutoff of 10 kDa and further subjected to gel filtration chromatography analysis using Superdex 200 increase 10/300 GL column (final buffer conditions-20 mM Tris, 150 mM NaCl). Purity and homogeneity of both the component (HlgA and HlgB) proteins of γ-HL was further monitored using SEC profile, and relevant peak fraction was monitored with 12% SDS-PAGE to assess the purity. Standard SDS-PAGE protocol was followed for the two protein components.

### Rabbit erythrocyte lysis assay

About 2-3 ml of whole blood was isolated from *Oryctolagus cuniculus* and was centrifuged at 3000 rpm for 5 minutes iteratively using 1x phosphate buffered saline (PBS) to selectively obtain erythrocytes for hemolysis assay with bicomponent HlgAB. A solution of 1 percent RBCs was finally prepared in 1x PBS buffer. 2 percent Tween-20 detergent was used as positive control to monitor change in absorbance. RBCs were treated with 1 μM HlgA and 1 μM HlgB and incubated in shaking conditions at 37°C for a duration of 30 minutes. Optical density was measured at 620 nm. Data points were recorded using Varioskan Flash microplate reader (Thermo Scientific). All experiments were performed in triplicates. Control and lysed RBCs were further visualized under room temperature NS-TEM. Rabbit whole blood was obtained upon approval of Institutional Animal Ethics Committee (IAEC), Indian Institute of Science, Bangalore (CAF/Ethics/881/2022).

### Preparation of Large, Unilamellar Vesicles (LUVs) of eggPC and cholesterol

For our study, LUVs constituted of eggPC and cholesterol in 1:1 molar ratio. Avanti Polar Lipids, L-α-phosphatidylcholine (Egg, Chicken) (egg PC) lipid and cholesterol were used for the preparation of the large unilamellar vesicles (LUVs) by lipid extrusion method. EggPC and cholesterol (in 1:1 molar ratio) were initially dissolved into 1 ml chloroform at room temperature followed by desiccation to remove the residual chloroform to get a thin lipid film. This pre-formed thin layer of egg PC lipid and cholesterol was further dissolved into 25 mM Tris, 150 mM NaCl buffer of pH 7.8 with gentle mixing and further incubated at 45°C (above the transition temperature) for 30 min. The resulting final lipid suspension in a colloidal state was subsequently processed by extrusion. For extrusion, we used 200 nm poresize polycarbonate membranes, and the colloidal lipid suspension was passed through the polycarbonate membrane for 8 times using Avanti Polar mini-extruder apparatus. Final extruded solution containing large unilamellar vesicles (LUVs) were stored at 4°C until further use.

### Generating oligomeric pore complexes of bicomponent HlgAB in 20 percent of 2-Methyl-2,4-pentanediol and eggPC-cholesterol LUVs

20 percent (v/v) 2-Methyl-2,4-pentanediol (MPD) was added to equimolar concentrations of HlgA and HlgB components and thoroughly mixed, followed by incubation at 37°C in water bath. The resultant pore complexes were loaded onto 10% SDS-PAGE without heat denaturation for the observation of SDS-stable oligomeric band and further observed under NS-TEM.

Equimolar HlgA and HlgB were further treated with freshly prepared eggPC-cholesterol LUVs in a protein:lipid ratio of 1:25, followed by incubation at 37°C in water bath. Untreated control LUVs and treated LUVs were monitored under NS-TEM for formation of oligomeric pore complexes.

### Sample preparation and data acquisition for room temperature negative staining TEM (NS-TEM)

3 μl aqueous sample of the protein after appropriate dilution was applied to a thin film of carbon coated copper grids (300 mesh size) which was glow discharged for 120 seconds before sample preparation. The sample was further stained with 2% uranyl acetate for 30 sec, followed by gentle blotting to remove excess stain from over the grid. These carbon-coated copper grids were then visualized under a room temperature 120 kV Talos L120C transmission electron microscope equipped with bottom-mounted Ceta camera (4000 × 4000 pixels) at magnification ranging between x57000-x92000.

### Image processing for NS-TEM micrographs

Different particle projections were manually picked using e2projectmanager.py (EMAN2.1)^52^, followed by extraction of particles from raw micrographs using e2boxer.py. ~6000-7000 particles for each dataset were further classified using reference-free 2D classifications in EMAN2.1, RELION2^53^ and SIMPLE^54^ programs.

### Sample preparation and data acquisition for single particle cryo-EM

GloQube glow discharge system was used to glow discharge Quantifoil R1.2/1.3 Cu 300 mesh holey carbon grids (Electron Microscopy Sciences) for 90 seconds at current of 20 mA prior to sample vitrification. About 3 μl of protein-treated LUVs sample (solution state) was applied onto the glow discharged grids and incubated for 10 seconds, followed by blotting and plunge-freezing (FEI Vitrobot Mark IV) into liquid ethane.

Data acquisition was performed using a 200 kV Talos Arctica (Thermo Scientific) equipped with K2 Summit Direct Electron Detector (Gatan, Inc.). About 2287 cryo-EM movies (20 frames/movie) were acquired with calibrated dosage of 2 e^-^/Å^2^/frame and a defocus range of −0.8 μm to −3 μm. Nominal magnification of x54000 was used for data acquisition, which corresponded to an Å/pixel value of 0.92. Automatic data collection was set up using LatitudeS data acquisition software (Gatan, Inc.).

### Data processing pipeline for cryo-EM single particle analysis

About 1,000,000 particles belonging to 2D lattice arrays were automatically picked using template picker and extracted with a box size of 400 pixels (Å/pixel= 0.92). Reference-free 2D classifications were used to remove junk particles and obtain class averages corresponding to different orientations of square lattice packing. An electron density map was also generated from a particular 2D class corresponding to top views with aligned particles using relion_reconstruct program in RELION3.1^53^.

For reconstruction of octahedral superassembly, manual picking and subsequent template-based automatic picking yielded 46595 particles from 2287 micrographs. With a box size of 400 pixels, these particles were extracted and classified rigorously to obtain 11634 choicest and well-centered particles. A subset of these particles was used to generate an *ab-initio* model without imposition of any symmetry, which revealed an octahedral arrangement of octameric pore complexes. Another initial model with octahedral symmetry illustrated higher resolution features throughout the map. This octahedral initial model was lowpass filtered to a resolution of 30 Å and used as the reference model for auto refinement of the abovementioned 11634 particles. CTF refinement followed by Bayesian polishing and one round of homogeneous refinement in CryoSPARC^55^ improved the final global resolution of the electron density map to 3.55 Å. The local resolution estimates were made using Blocres^56^.

### Atomic model building and model validation

Rigid-body fitting of the available PDB model for LukF-Hlg2 hetero-octamer (PDBID: 3B07) was performed in UCSF Chimera^13^. PHENIX’s phenix.dock_in_map and phenix.real_space_refine tools were used to refine the initial model 3B07^57^. The refined model was imported in COOT^58^ to manually correct the side chain fitting and annotate additional densities to previously missing residues, including the N-terminal domain of HlgA. This was followed by another round of real space refinement of the entire atomic model using phenix.real_space_refine. Final model validation statistics were generated using cryo-EM comprehensive validation and EMRinger tools in PHENIX^57,59^. Molecular interactions were plotted using PDBsum and visualizations were performed using PyMOL and UCSF ChimeraX^60^. Model refinement and validation statistics are also provided separately in Table 1.

## ACKNOWLEDGEMENTS

We acknowledge Department of Biotechnology, Department of Science and Technology (DST) and Science, and Ministry of Human Resource Development (MHRD), India for funding and cryo-EM facility at IISc-Bangalore. We acknowledge DBT-BUILDER Program (BT/INF/22/SP22844/2017) and DST-FIST (SR/FST/LSII-039/2015) for National Cryo-EM facility at IISc, Bangalore. We acknowledge the financial support from the Ministry of Human Resource Development (MHRD) (Grant Number-STARS-1/171). We acknowledge DBT-IISc partnership program phase II for the negative staining TEM facility at Biological Sciences Division, IISc and cryo-EM data processing cluster. We acknowledge the high-performance computing cluster “Beagle” at Biological Sciences Division, IISc. We also thank Mr. Clayton Fernando for preparing cryo-EM grids and aiding cryo-EM data collection.

## AUTHOR INFORMATION

### Affiliations

Molecular Biophysics Unit, Indian Institute of Science, Bangalore-560012, Karnataka, India Suman Mishra, Anupam Roy and Somnath Dutta

### Contributions

S.M. and A.R. purified recombinant protein, performed biophysical characterization and negative staining TEM studies. S.M. performed NS-TEM and cryo-EM image processing. S.M. and A.R. performed atomic model building. S.M., A.R. and S.D. analyzed the data, interpreted results, and wrote the manuscript. S.M. and A.R. prepared the figures. S.D., S.M. and A.R. designed the research. S.D. provided funding.

### Corresponding author

Correspondence to Dr. Somnath Dutta.

### ETHICS DECLARATIONS

The authors declare no competing interests.

