## Supplemental Figure for "Cryo-EM-based structural insights into supramolecular assemblies of γ-Hemolysin from *Staphylococcus aureus* reveal the pore formation mechanism"

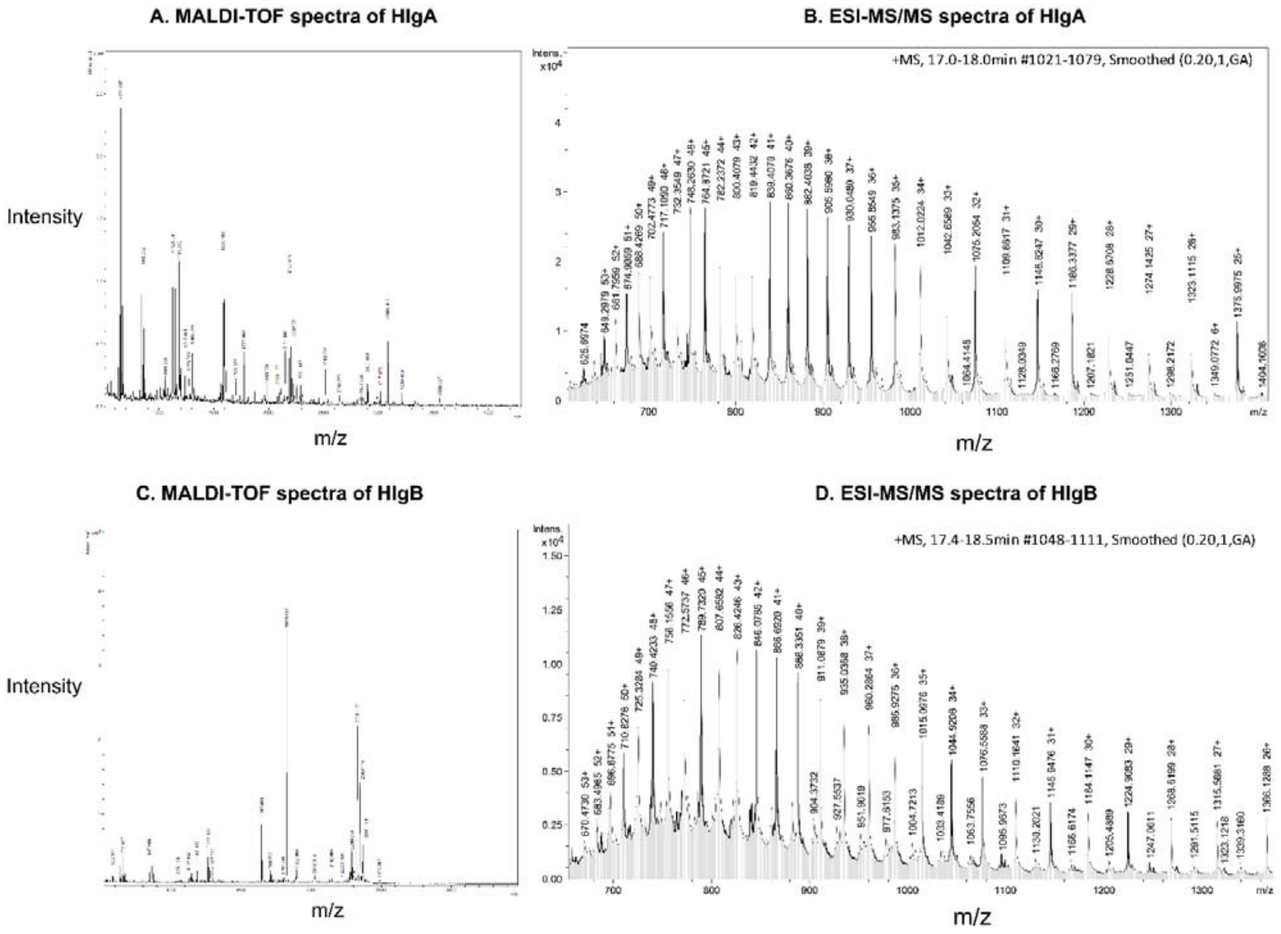


**Figure S1. Mass spectrometry-based characterization of the toxin components, HlgA and HlgB. A, C.** MALDI-TOF spectra of trypsin-digested HlgA and HlgB, respectively (intensity vs. m/z). **B, D.** ESI-MS/MS spectra of HlgA and HlgB, respectively (intensity vs. m/z).

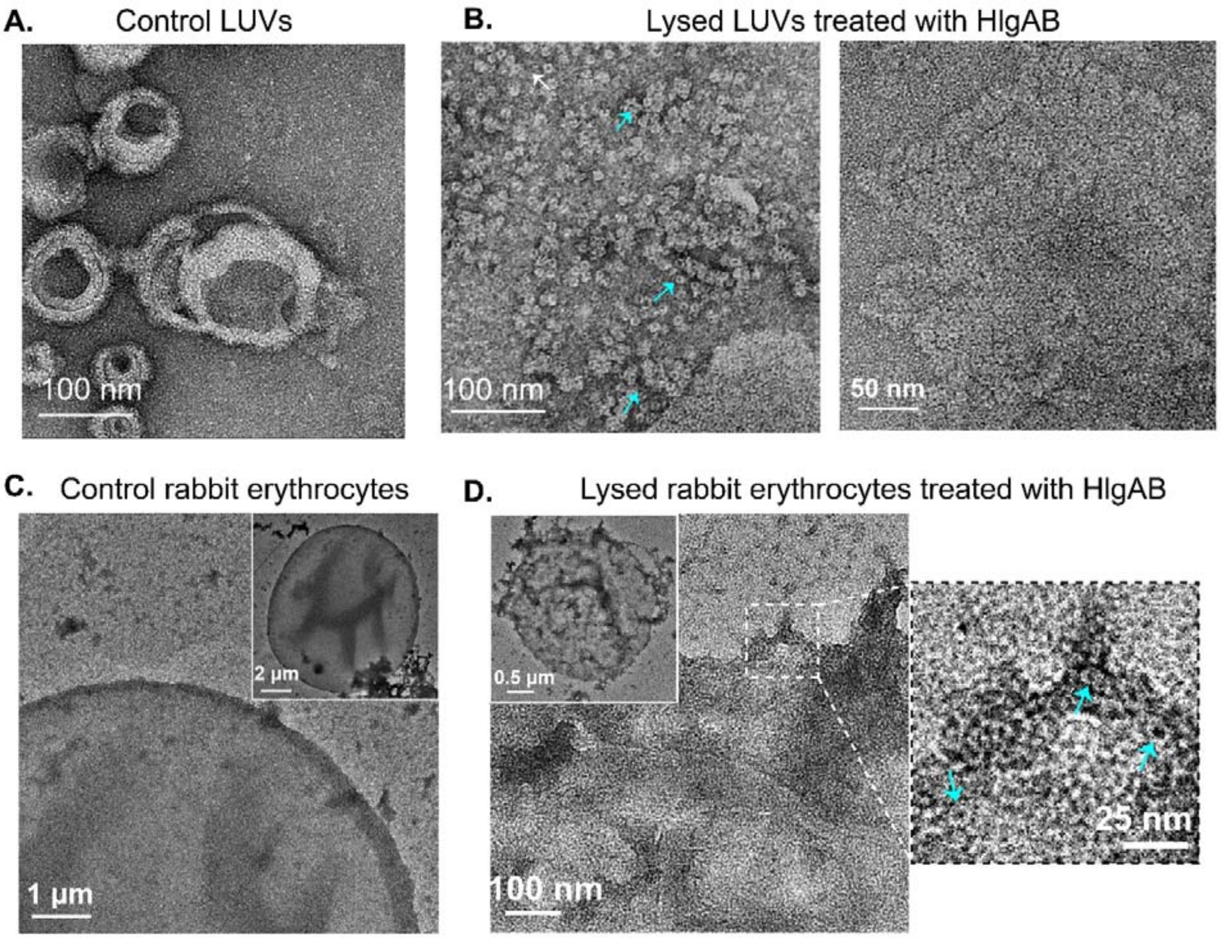


**Figure S2. Gallery of NS-TEM micrographs of HlgAB octameric pore complex induced by EggPC-Cholesterol LUVs and rabbit erythrocytes. A.** Representative NS-TEM micrograph of large unilamellar vesicles (LUVs) composed of equimolar concentrations of eggPC and cholesterol, as negative control. **B.** EggPC-cholesterol LUVs incubated with equimolar concentrations of HlgA and HlgB at 37°C and observed under NS-TEM. Corresponding micrographs demonstrate the rupturing of LUVs (left) and formation of HlgAB pore complexes. Blue arrows indicate several clusters of HlgAB oligomers. 2D lattice arrays of HlgAB pore complexes were also observed (right). **C-D.** Representative NS-TEM micrographs of untreated (C) and HlgA/HlgB treated (D) rabbit erythrocytes. Untreated erythrocytes were observed for membrane integrity. Inset shows the complete intact erythrocyte for reference at lower magnification. Treated erythrocytes illustrated ruptured erythrocyte membrane. Enlarged view of a selected area is shown (extreme right) with pore complexes indicated with blue arrows. Inset represents the complete treated erythrocyte for reference at lower magnification. Enlarged view of micrographs in insets are shown separately in Figure S4.

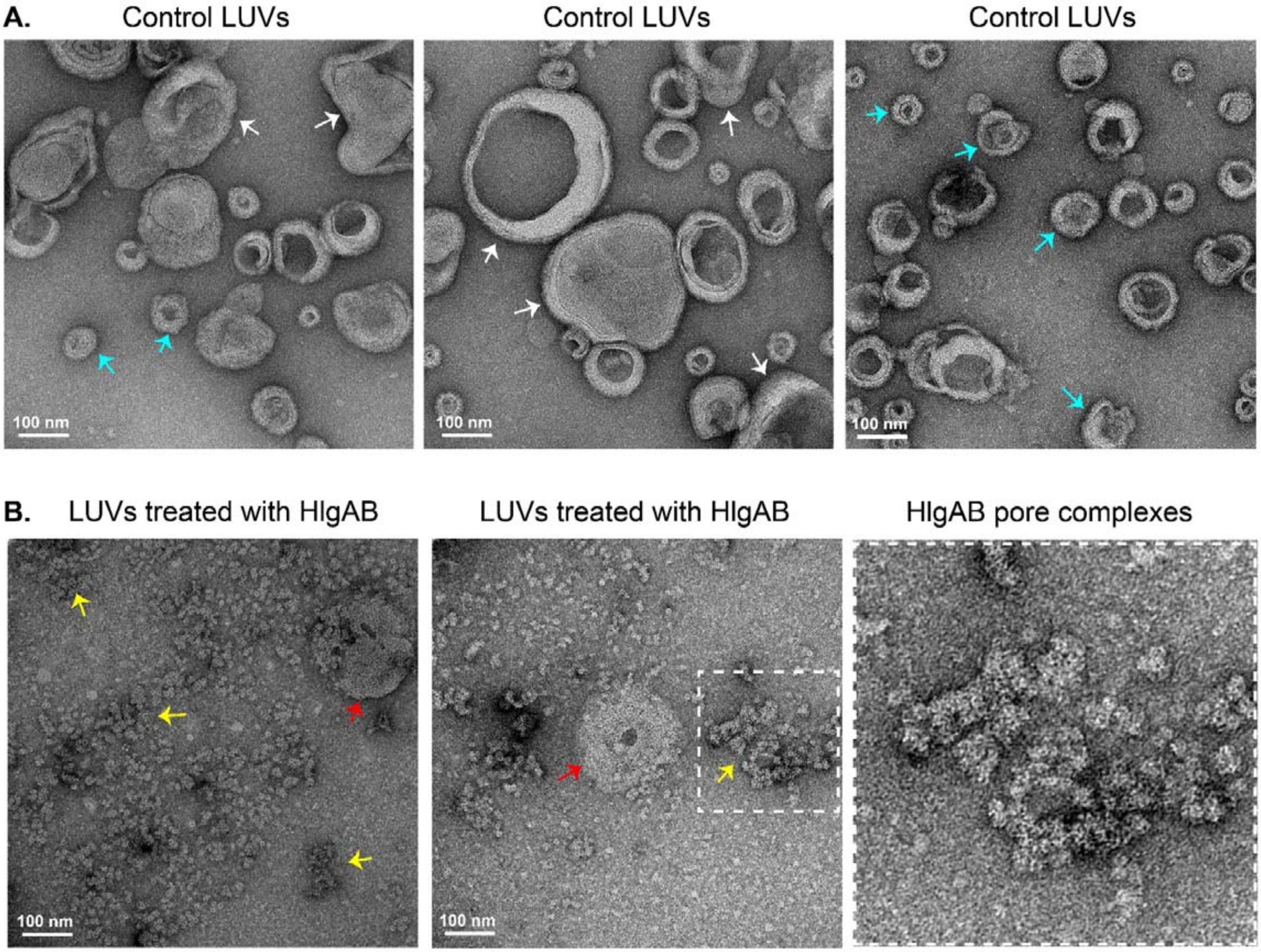


**Figure S3. Gallery of NS-TEM micrographs of negative control and toxin (HlgA, HlgB) treated LUVs. A.** LUVs comprising of equimolar concentrations of eggPC, and cholesterol were used as a model membrane for this study. Untreated LUVs were observed to be spherical under NS-TEM. White and cyan arrows indicate large and small liposomes, respectively. All liposomes were observed to be globular and intact. **B.** LUVs treated with equimolar concentrations of HlgA and HlgB components and incubated at 37°C. Rupturing of liposomes and clusters of protein complexes in the background were apparent. Red and yellow arrows indicate rupturing liposomes and clusters of HlgAB pore complexes, respectively. One such cluster is highlighted in dashed square and an enlarged view (bottom right) is presented for visualization.

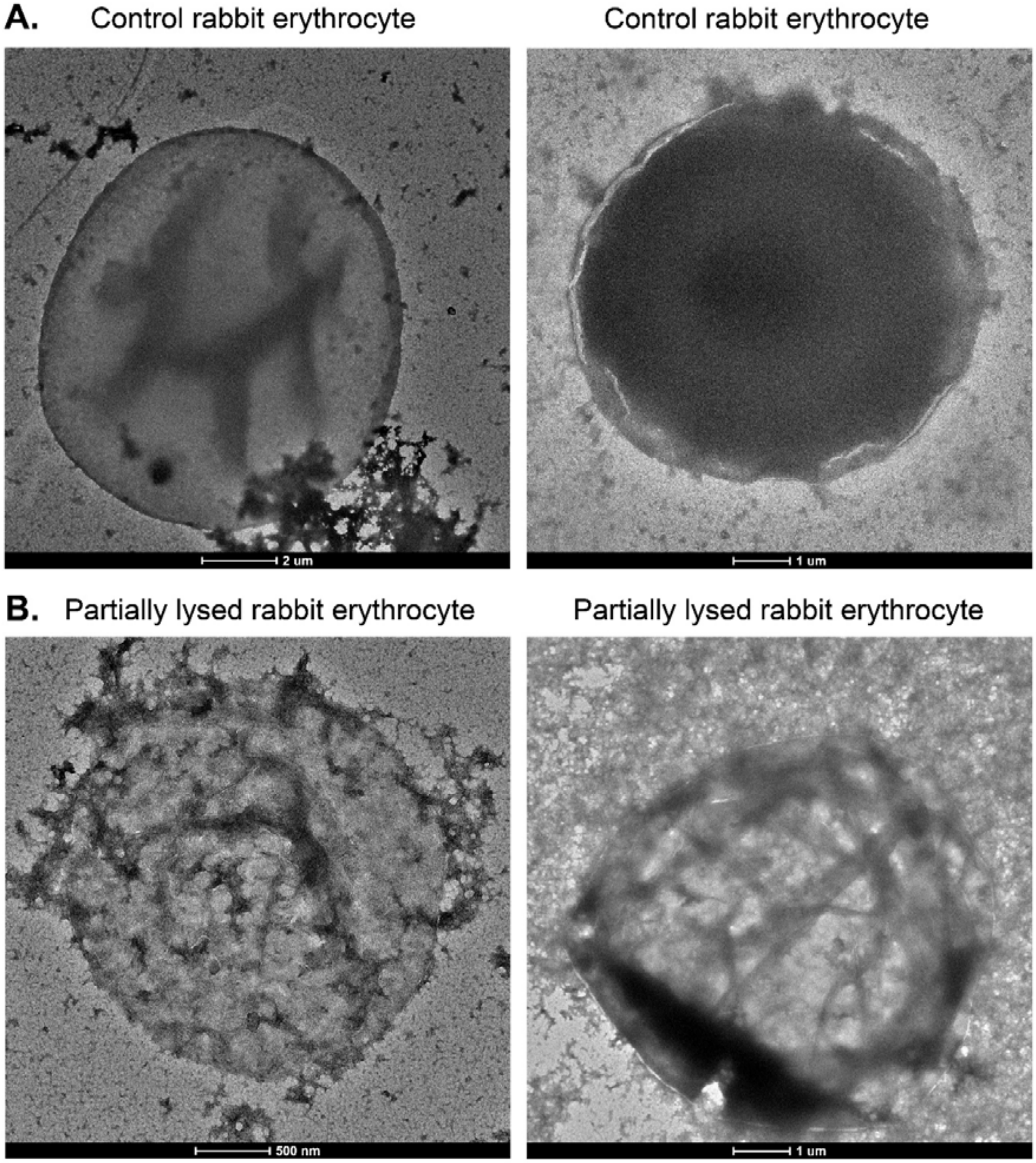


**Figure S4. Untreated and partially lysed rabbit erythrocytes upon treatment with γ-HL.** **A.** Representative NS-TEM micrographs of intact rabbit erythrocytes. RBCs appear to be globular and retain their membrane integrity and curvature. **B.** The erythrocytes were treated with HlgA and HlgB components (1:1 molar ratio) and incubated for 30 minutes before imaging under NS-TEM. Representative micrographs illustrate perturbed and partially lysed erythrocytic membranes. An enlarged view of intact and partially lysed erythrocytes present in insets of Figure S2C-D are also shown here for reference.

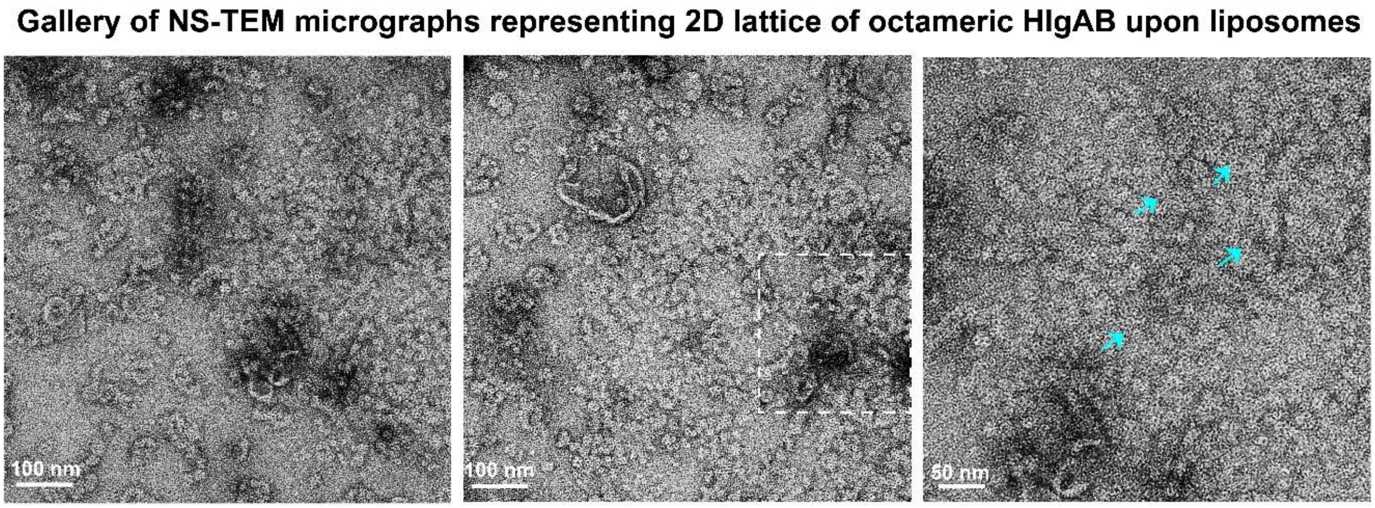


**Figure S5. Gallery of NS-TEM micrographs representing 2D lattice of octameric HlgAB upon liposomes.** 2D lattice arrays of HlgAB pore complexes were observed upon EggPC-Cholesterol liposomes (left and centre). An enlarged view of one such lattice (enclosed in white dotted rectangle) is presented (right), indicating HlgAB individual pore complexes with cyan arrows.

### Cryo-EM gallery of HlgAB oligomers with EggPC-Cholesterol liposomes

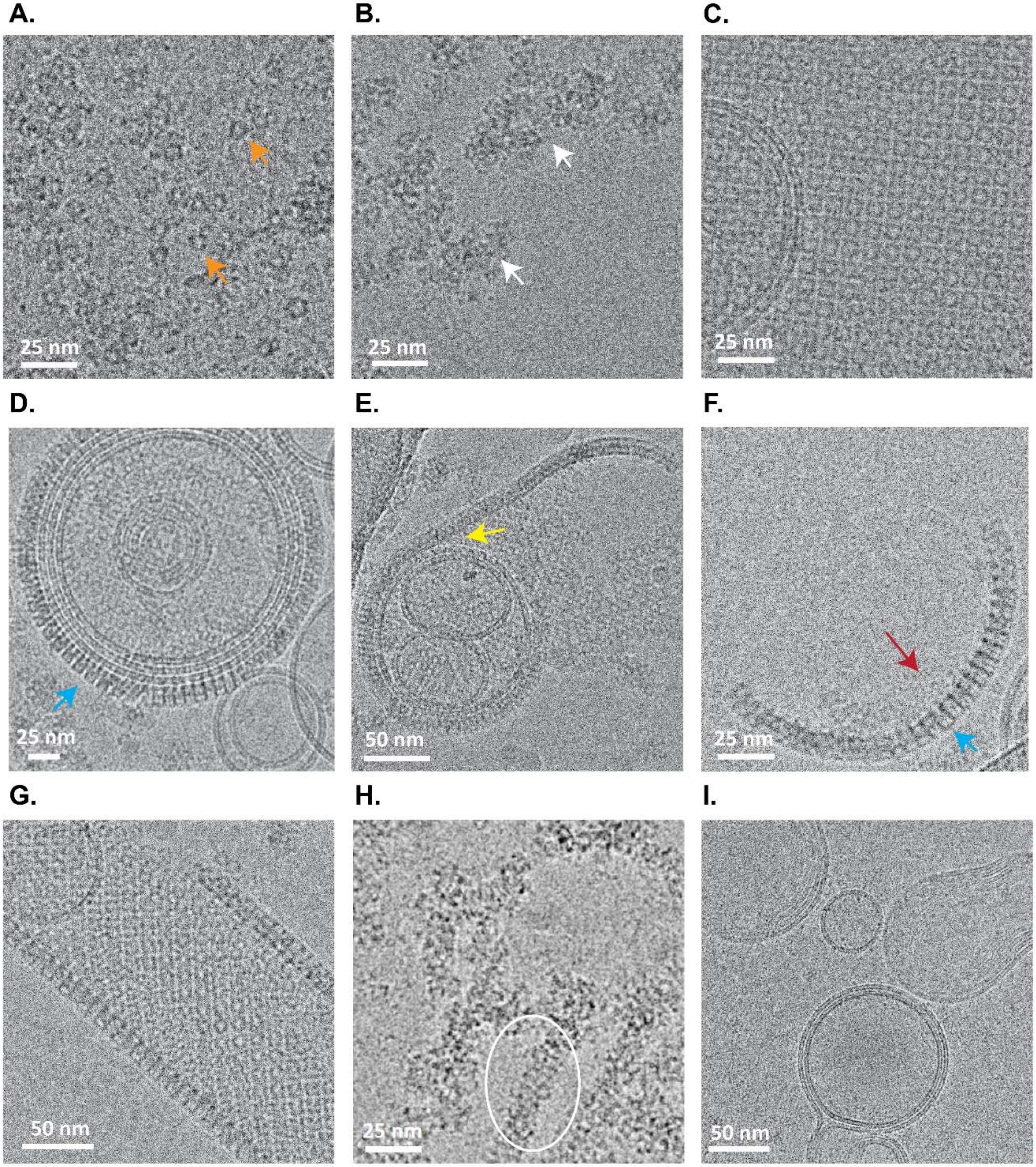

**Figure S6. Cryo-EM gallery of HlgAB oligomers with EggPC-Cholesterol liposomes. A.** Top views of individual HlgAB pore complexes, highlighted with orange arrows. **B.** Two non-specific clustering of pore complexes are indicated with white arrows. **C.** Micrograph with 2D lattice of HlgAB pore complexes uniformly extending in all directions. **D.** A representative micrograph showing 2D lattice formation curving upon a spherical and intact LUV (highlighted in blue arrow). **E.** A representative micrograph illustrating a layer of HlgAB lattice array peeling off a multilamellar vesicle (marked in yellow arrow). **F.** Side view of a peeled arc of HlgAB lattice. Blue and red arrows indicate convex and concave faces of the arc, respectively. **G.** A representative micrograph showing 2D lattice arrangement of HlgAB pore complexes over an elongated curved membrane surface. **H.** Several clusters of HlgAB oligomeric pores can be observed. One such linear cluster of matured pores is encircled in a white solid ellipse. Scale bar corresponds to 25 nm. **I.** Control untreated EggPC-Cholesterol LUVs for reference.

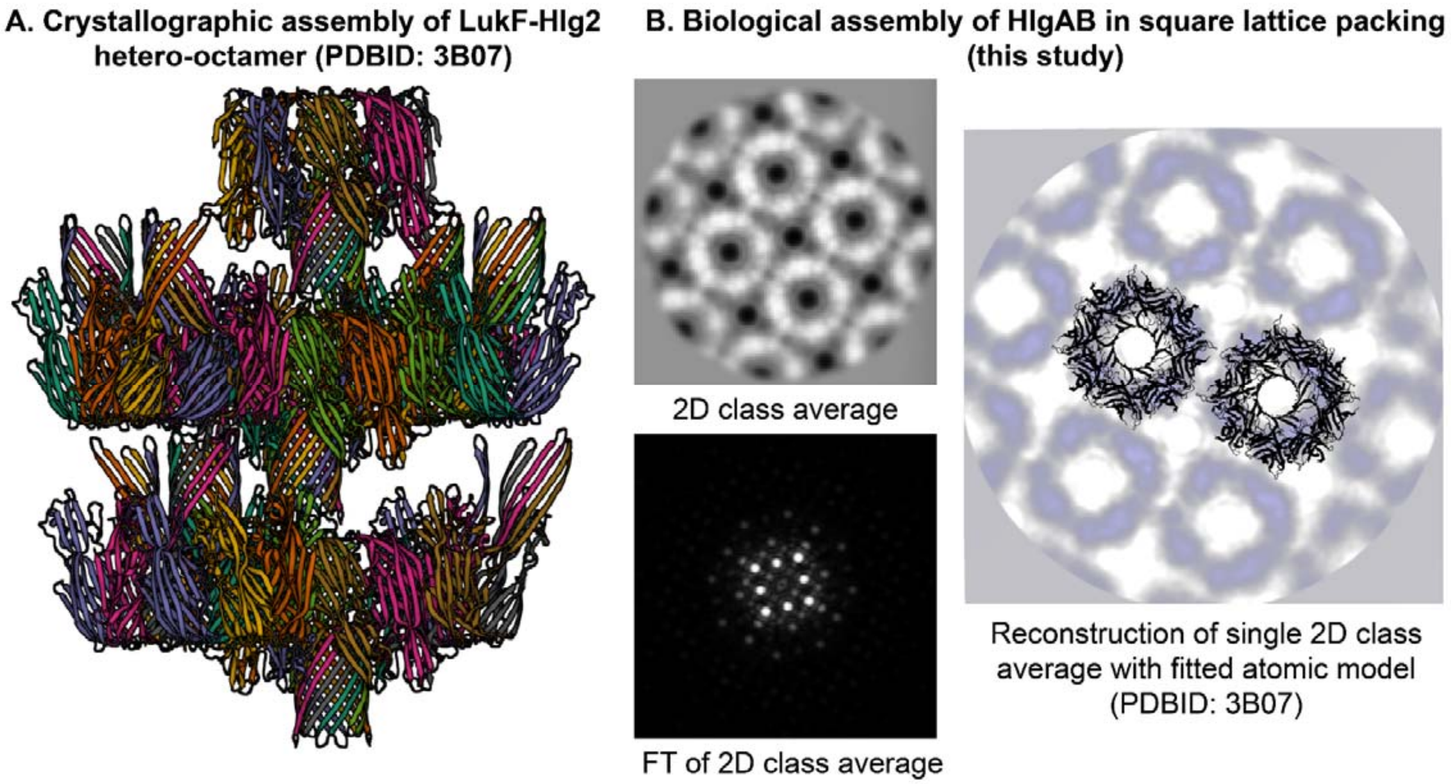


**Figure S7. Comparison between the crystallographic assembly and biological assembly of bicomponent HlgAB. A.** Crystallographic assembly of LukF-Hlg2 hetero-octamer (PDBID: 3B07) reveals antiparallel arrangement of octameric pore complexes within a plane. **B.** 2D class average of square lattice arrangement of bicomponent HlgAB (top left), FT of 2D class average (bottom left), and the rigid body fitting of octameric pore complexes (PDBID: 3B07) in corresponding reconstruction of single 2D class average using relion_reconstruct (right) reveal biological assembly of octameric pores in a parallel arrangement in square lattice.

**A. Gallery of NS-TEM micrographs of octahedral superassembly induced by lipid membranes**

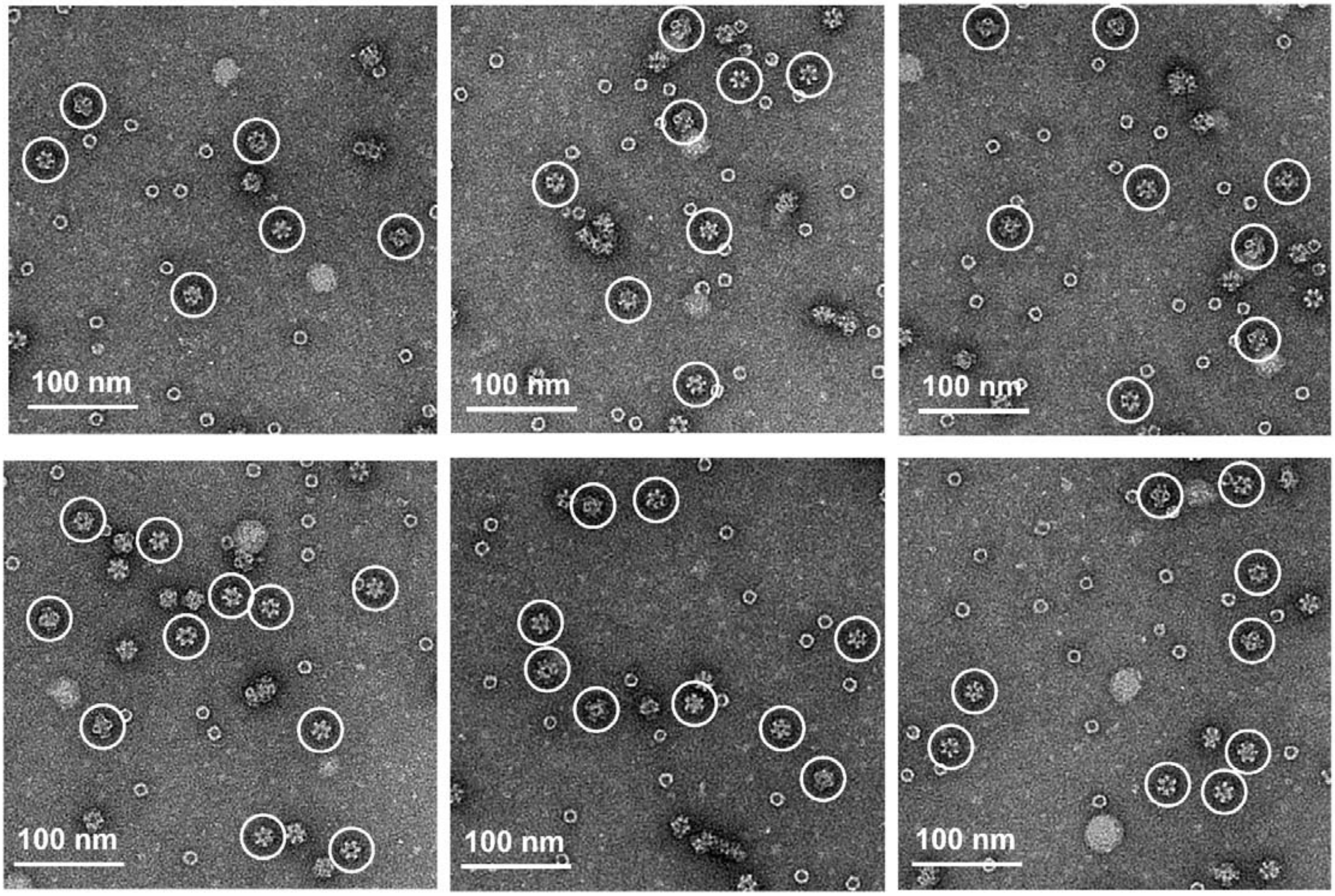


**B. Representative NS-TEM 2D class averages of supramolecular assemblies of HlgAB**

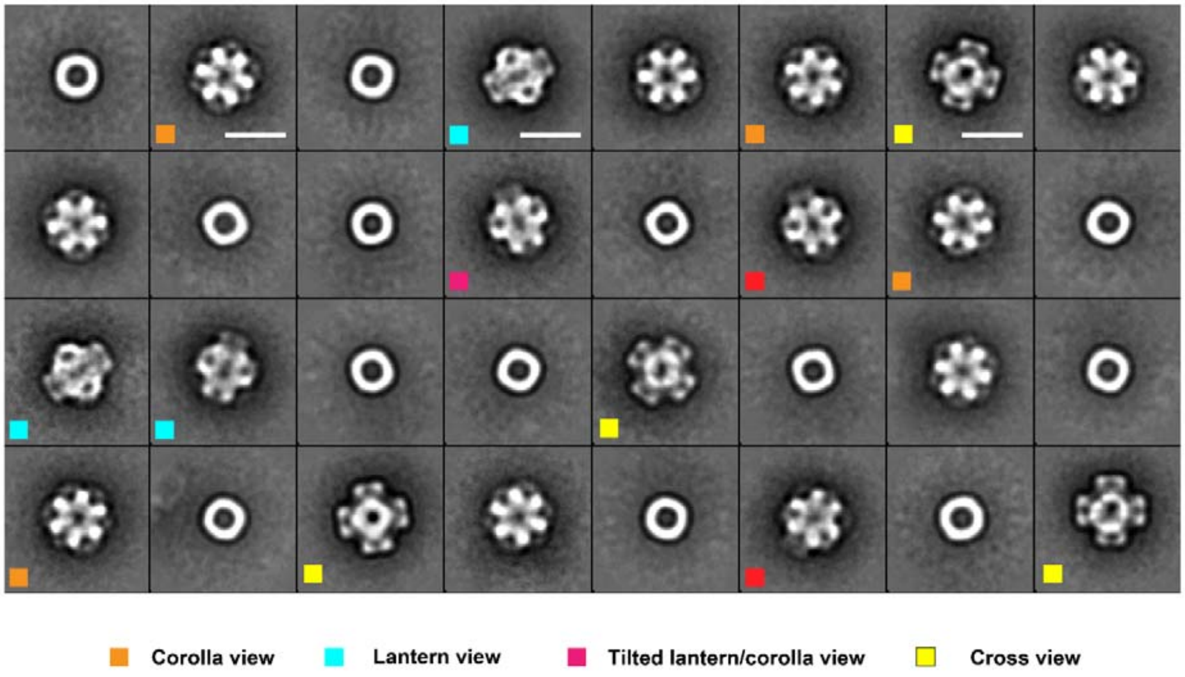

**Figure S8. Gallery of NS-TEM micrographs of octahedral super assembly induced by lipid membranes and their respective 2D class averages. A.** In addition to individual HlgAB octamers, clusters of HlgAB pore complexes and lattice arrays, another predominant species formed upon incubation of the two components with EggPC-Cholesterol liposomes, was the octahedral supramolecular assembly of octameric HlgAB pore complexes. Six representative micrographs illustrate the monodispersed distribution of these super assemblies (encircled in white) under room temperature NS-TEM. **B.** Representative NS-TEM 2D class averages of supramolecular assemblies of HlgAB observed in addition to the clustering and square lattice packing. Corolla view (marked with orange), cross view (marked with yellow), lantern view (marked with cyan), and tilted lantern/corolla view (marked with red) of the superassembly particles indicate uniform angular sampling. Scale bar denotes 20 nm.

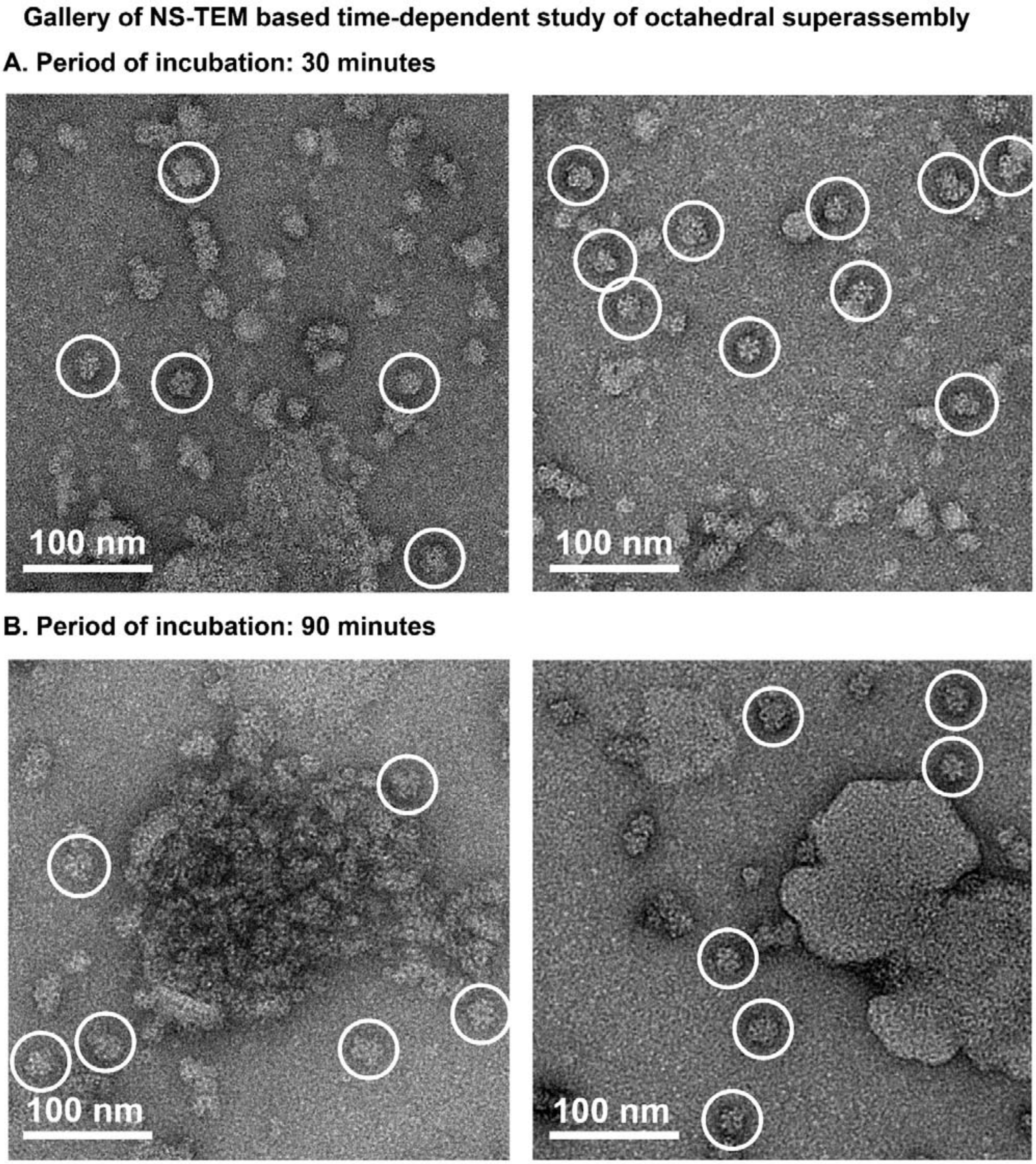


**Figure S9. Gallery of NS-TEM based time-dependent study of formation of octahedral super assembly.** In a time-dependent study of bicomponent HlgAB with EggPC-Cholesterol liposomes, population of octahedral HlgAB supramolecular assembly was observed in **A.** 30 minutes, **B.** 90 minutes post incubation at 37°C under NS-TEM. Such species are encircled in white solid lines.

Representative NS-TEM micrograph and corresponding 2D class averages of octahedral superassembly of octameric HlgAB induced in the presence of 20% 2-Methyl-2,4-pentanediol

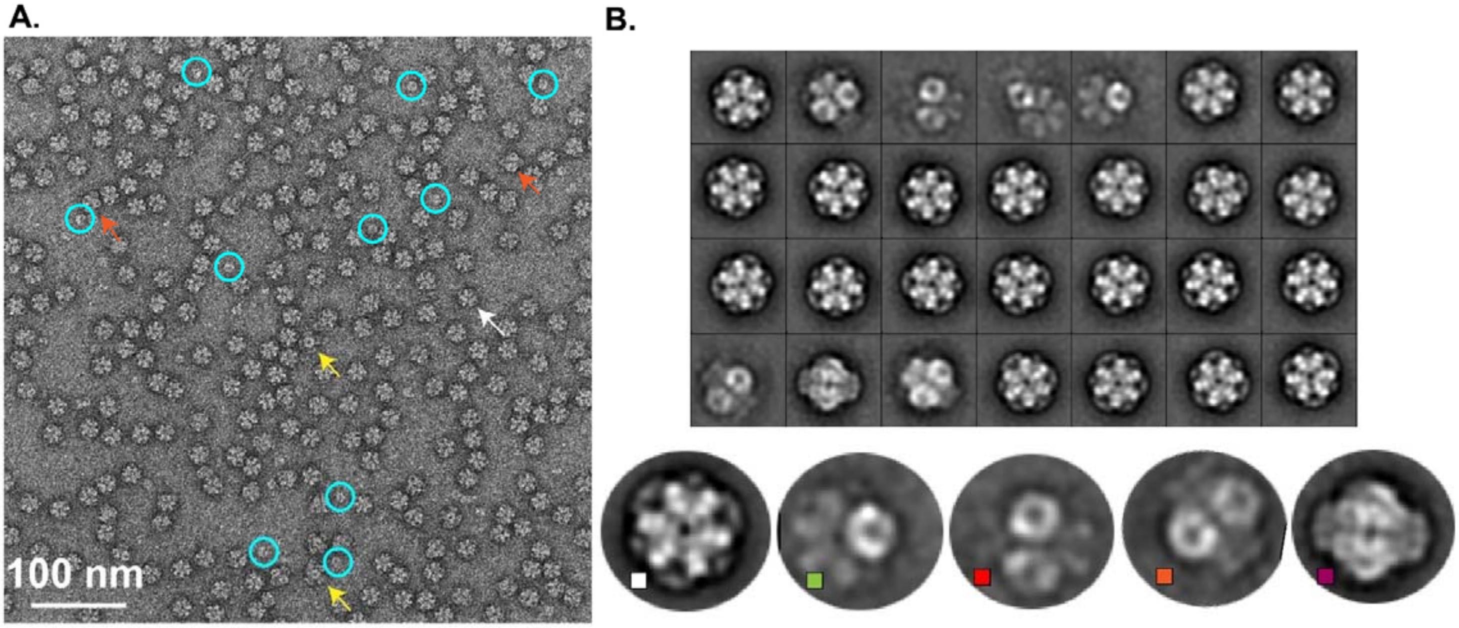


**Figure S10. Octameric assembly and octahedral super assembly of bicomponent HlgAB induced by 20% 2-Methyl-2,4-pentanediol. A.** Representative raw micrograph (left) with octameric pore complexes of HlgAB upon treatment with 20% MPD and incubation at 37°C for 30 minutes. Individual pore complexes are marked with white arrows. Raw particles and corresponding 2D class averages show homogeneity in particle size distribution. Only top views of octameric pore complexes were visible under NS-TEM (class averages marked with white square). **B.** Octahedral supramolecular assemblies were also observed under NS-TEM (left) after overnight incubation at 37°C. Yellow, orange and white arrows in the micrograph indicate “cross” view, “lantern” view, and “corolla” view, respectively. Extended 2D classification with ~7300 particles (top right) and selected enlarged 2D class averages representing different views (colour coded accordingly) are also provided. These 2D classes agree with classification results in a membrane environment (refer Figure S8).

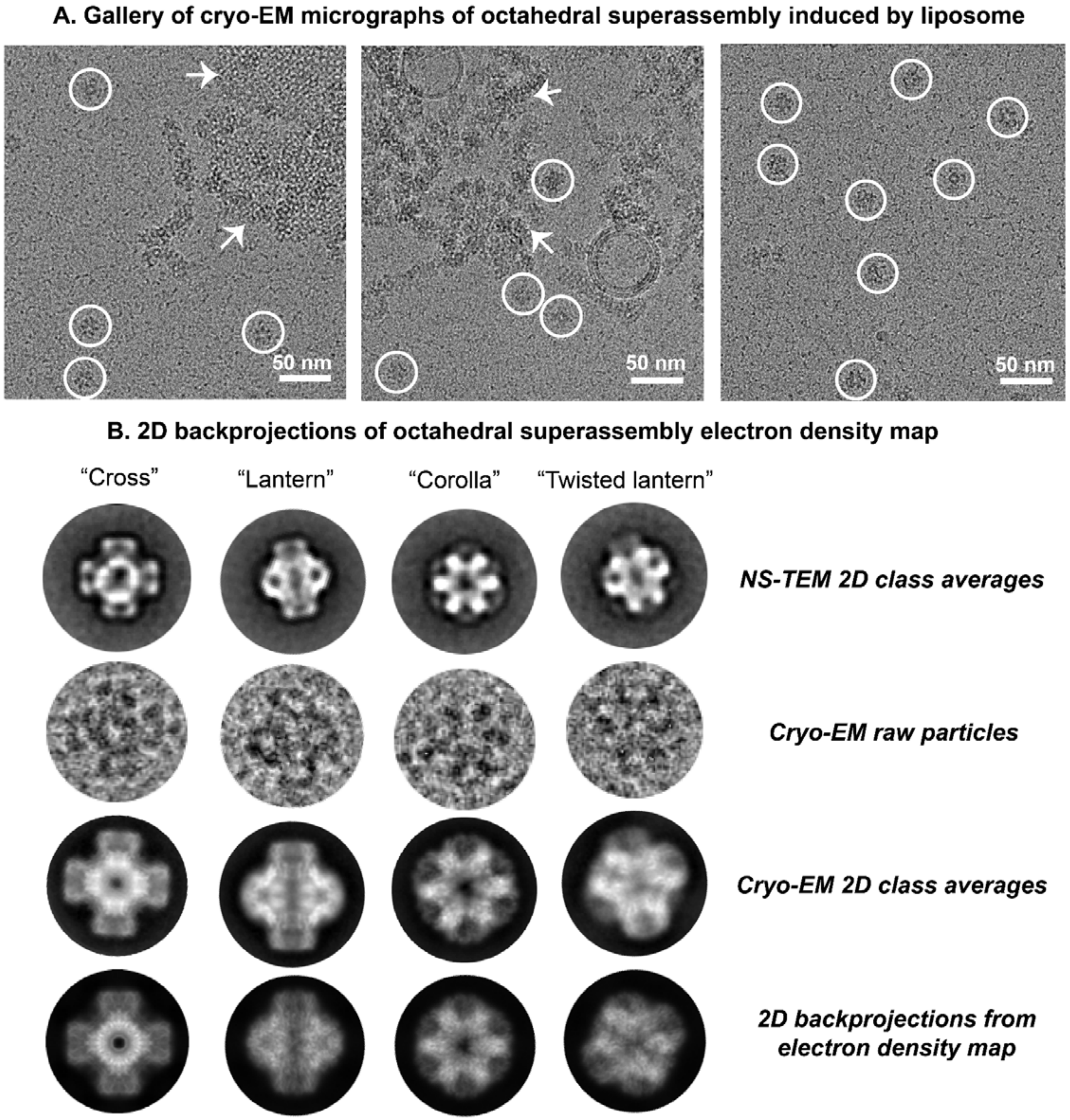
A. Gallery of cryo-EM micrographs of octahedral superassembly induced by liposome
50 nm
50 nm
50 nm
B. 2D backprojections of octahedral superassembly electron density map
"Cross"
"Lantern"
"Corolla"
"Twisted lantern"
NS-TEM 2D class averages
Cryo-EM raw particles
Cryo-EM 2D class averages
2D backprojections from electron density map

**Figure S11. Cryo-EM structural characterization of octahedral supramolecular assemblies of HlgAB. A.** A gallery of micrographs has been provided to illustrate the distribution of raw particles under cryo-EM, further used for 3D reconstruction of the octahedral super assembly of HlgAB using single particle analysis. Superassembly particles are encircled in white. Lattice of oligomeric HlgAB is denoted with white arrows. **B.** 2D back projections of the 3D reconstructed superassembly electron density map illustrate different orientations of the superassembly (cross, lantern, corolla, and twisted lantern), correlating with cryo-EM raw particles, NS-TEM, and cryo-EM 2D class averages. High resolution features of the electron density map (at a global resolution of 3.5 Å) are also evident from the back projections.

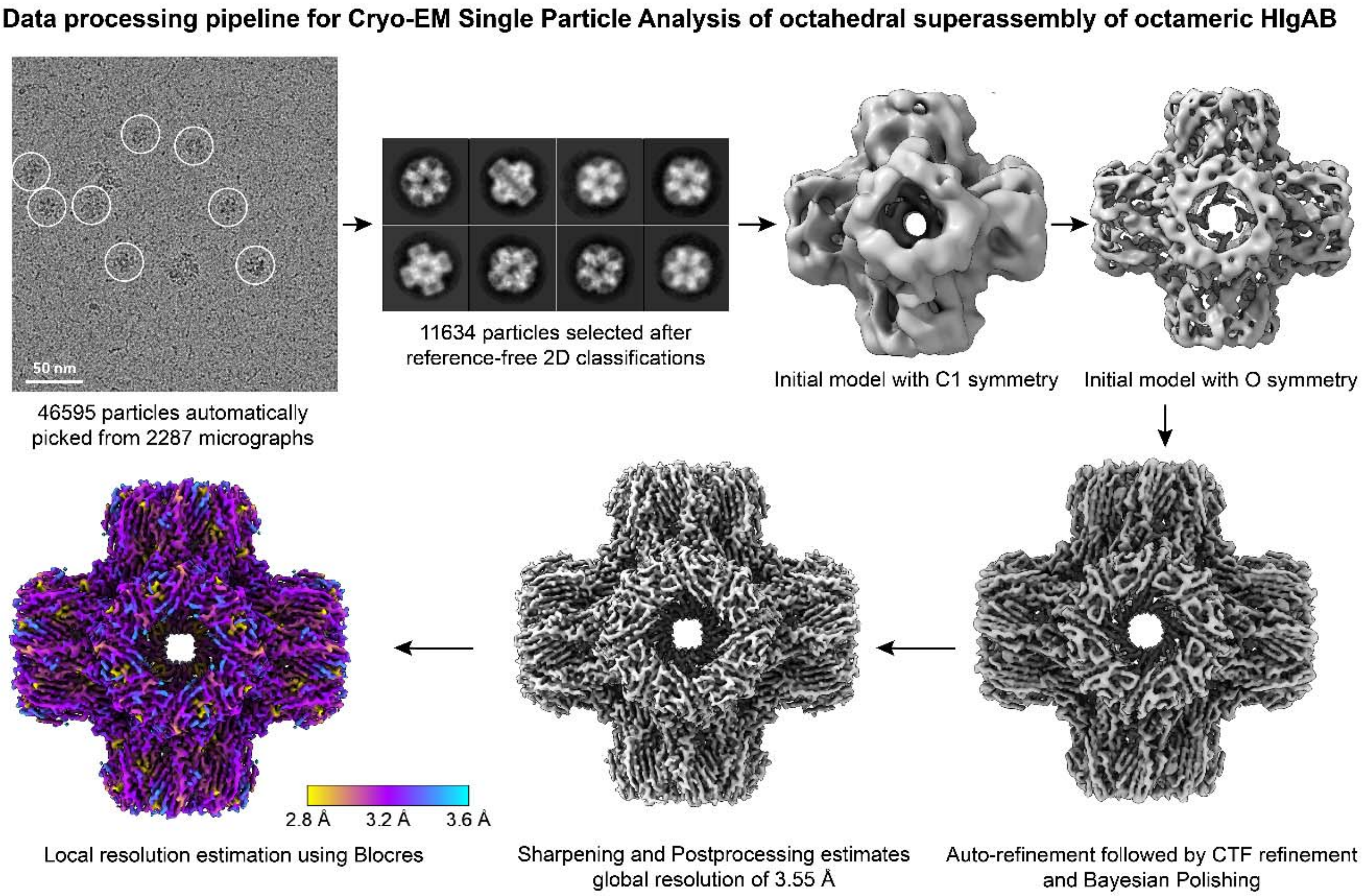


**Figure S12. Data processing pipeline for cryo-EM single particle analysis of octahedral superassembly of octameric HlgAB.** Initially, particles corresponding to octahedral superassembly were picked and classified to perform template-based automatic picking from 2287 micrographs. A total of ~46595 particles were extracted and further classified rigorously to obtain 11634 choicest and well-centred particles. An initial model without imposition of any symmetry was generated, which illustrated that octameric HlgAB pore complexes were arranged rim-to-rim in an octahedral fashion. Another initial model was generated with octahedral symmetry imposed, which further improved the features of the model. This model was lowpass filtered to 30 Å and used as a reference density map for auto-refinement of 11634 particles. CTF refinement, Bayesian polishing in RELION and subsequent refinement in CryoSPARC yielded an electron density map with gold standard FSC resolution of 3.55 Å. Local resolution estimates for the final electron density map using Blocres peaked at resolution of ~3.2 Å.

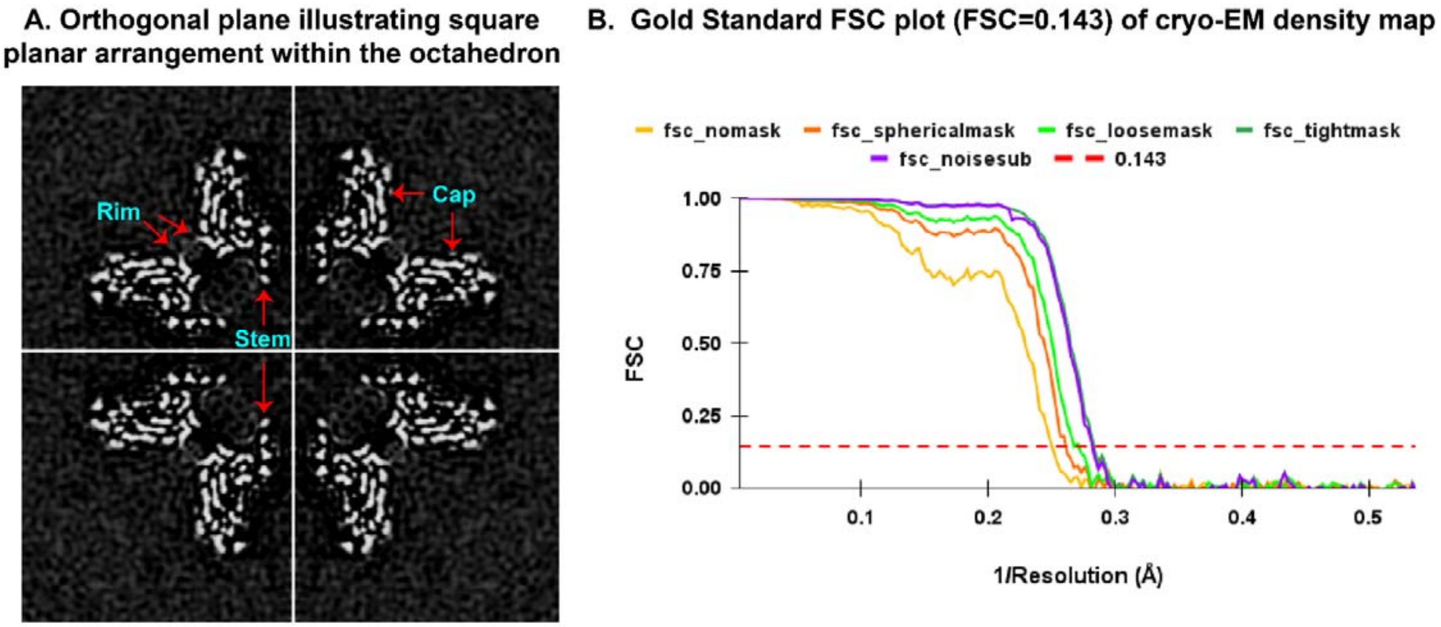


**Figure S13. Orthogonal plane and gold standard FSC for octahedral super assembly of γ-HL. A.** Orthogonal plane illustrating square planar arrangement within the octahedral supramolecular assembly. The arrangement of cap, rim and stem domains have been indicated with red arrows. **B.** Gold standard Fourier Shell Correlation (GS-FSC) predicts the global resolution of electron density map at 3.55 Å. Plot was generated for FSC vs. 1/Resolution (Å). Colour key for the curves has been provided.

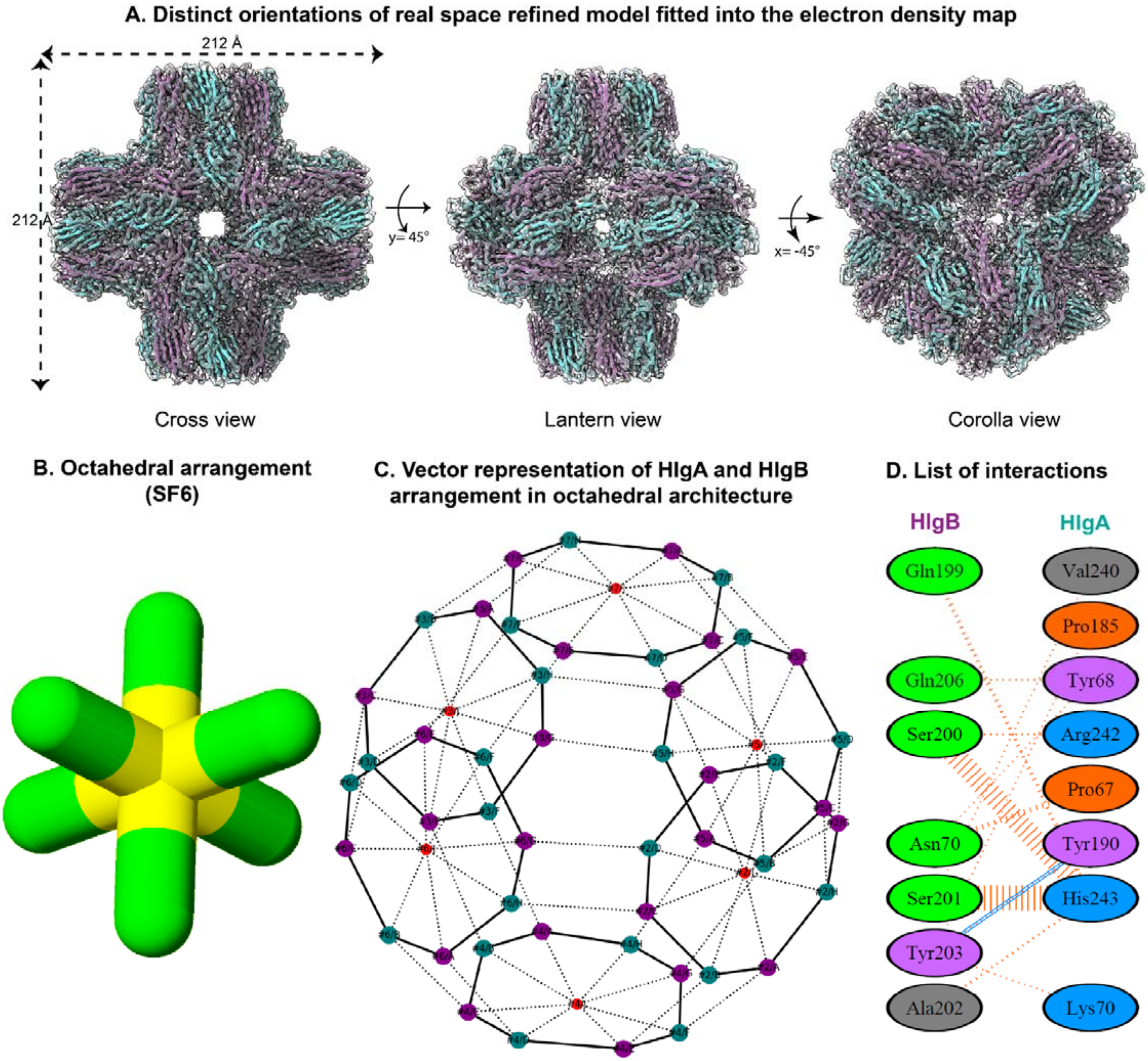
A. Distinct orientations of real space refined model fitted into the electron density map
212 Å
212 Å
y= 45°
x= -45°
Cross view
Lantern view
Corolla view
B. Octahedral arrangement (SF6)
C. Vector representation of HlgA and HlgB arrangement in octahedral architecture
D. List of interactions
HlgB
HlgA
Gln199
Val240
Pro185
Gln206
Tyr68
Ser200
Arg242
Pro67
Asn70
Tyr190
Ser201
His243
Tyr203
Ala202
Lys70

**Figure S14. Distinct orientations of atomic model fitted into the electron density map; insights into interactions between protomeric HlgA and HlgB into octameric assembly and octahedral superassembly. A.** Overall fitting of the atomic model for octahedral superassembly of bicomponent γ-HL from this study has been shown within transparent grey electron density map in three distinct orientations- cross view, lantern view and corolla view. HlgA and HlgB are marked cyan and purple, respectively for visualization. The diameter of the octahedral superassembly is ~212 Å. **B.** Octahedral arrangement in sulfur hexafluoride ($SF_6$). **C.** Vector representation of HlgA and HlgB arrangement in octahedral cage architecture. HlgA and HlgB are indicated in cyan and purple colour, respectively. Solid lines indicate six octameric assemblies of HlgAB. Dotted lines between octamers indicate inter-octameric contacts involved in the octahedron superassembly. **D.** Hydrophobic zipping contributed by the rim domain of both components HlgA and HlgB stabilizes the HlgAB superassembly interface.

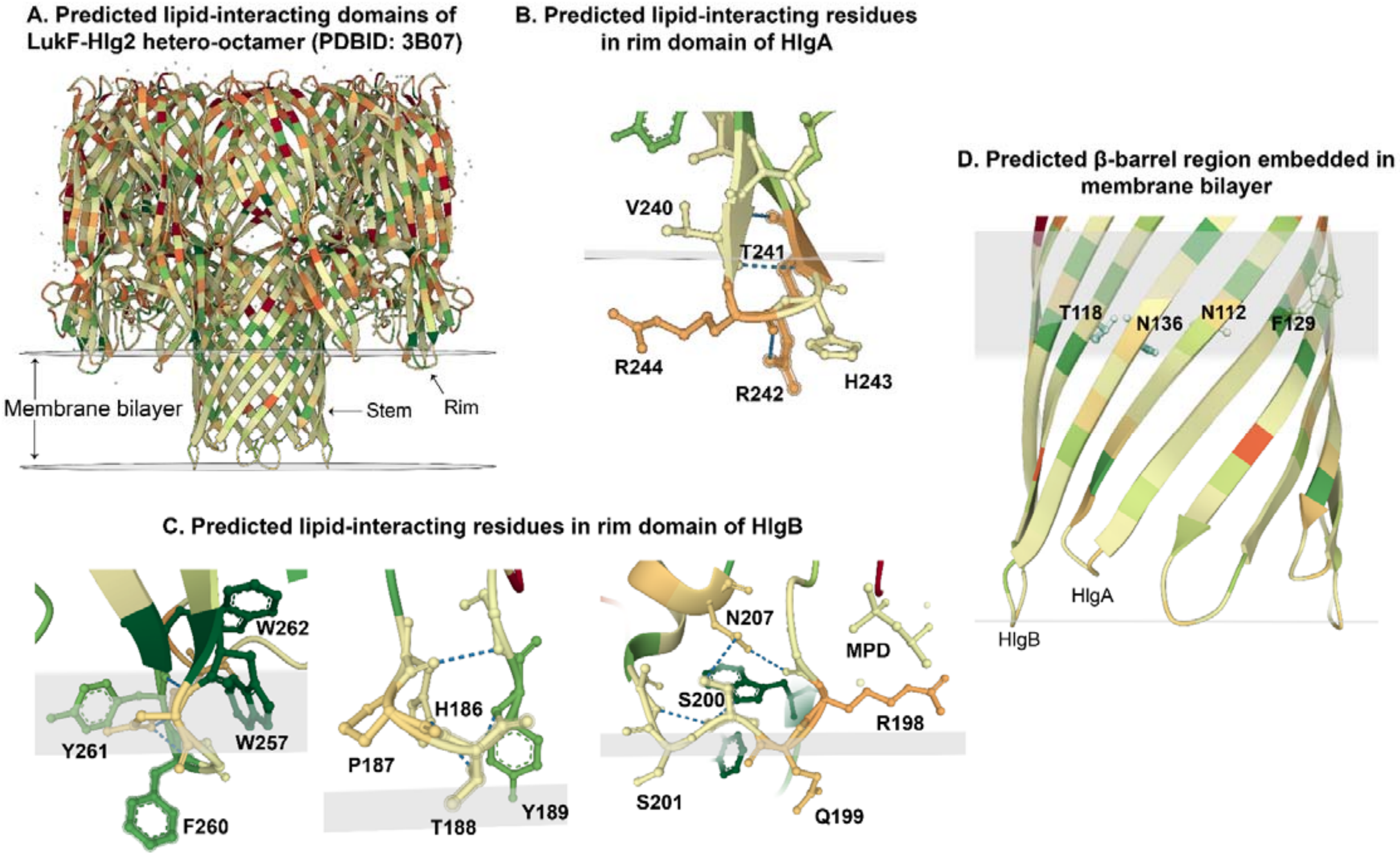


**Figure S15. Predicted lipid-interacting residues for the components LukF (HlgA) and Hlg2 (HlgB) for hetero-octameric pore complex (PDBID: 3B07). A.** Predicted lipid-interacting domains. Membrane bilayer and stem/rim domains are indicated. **B.** Predicted residues V240-R244 involved in lipid interaction in rim domain of HlgA. Lipid densities were observed at superassembly interface for these residues. Please refer Figure 5A. **C.** Predicted residues H186-Y189, R198-N207, W257-W262 involved in lipid interaction in rim domain of HlgB. Please refer Figure 5A. **D.** Predicted transmembrane β-barrel region consists of residues N112-F129 for HlgA and T118-T137 for HlgB. Densities for these residues were absent in the electron density map, suggesting disorder in absence of lipid environment. Please refer Figure S20.

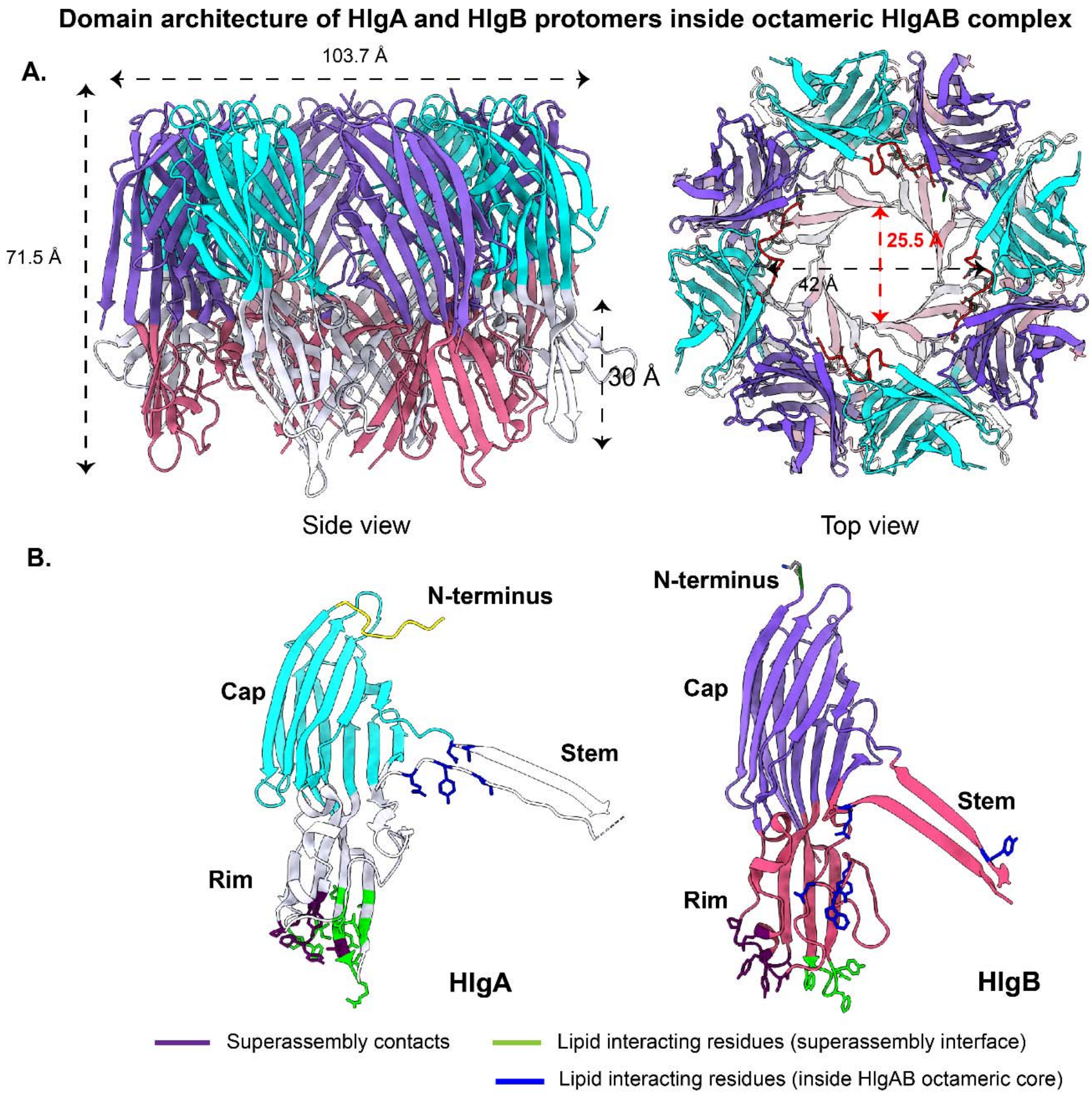


**Figure S16. Octameric topology and domain architectures of HlgA and HlgB protomers in octameric HlgAB pore complex. A.** A cartoon representation of HlgAB octameric pore complex from side view and top view respectively. The four protomeric subunits of HlgA and HlgB are shown in different domains colour compositions arranged in an alternative arrangement where N-terminal amino latch (shown in red colour) from four protomeric HlgA are spanning symmetrically (C4 rotational) through the octameric pore. The inner and outer diameter of octameric pore composed of cap domains are 42 Å & 103 Å, while the pore diameter and height of transmembrane β-barrel containing four β-hairpin segments of HlgA and HlgB are 25.5 Å and 30 Å, respectively. The overall height of the octameric pore complex is 71.5 Å. The pore complex has been shown to maintain a C4 rotational symmetry (pseudo C8) along the central pore axis. The C4 rotational symmetry is maintained by both cap as well as rim domains of HlgAB hetero octamers. **B.** Distribution of the critical lipid binding residues/elements and super-assembly governing residues located in rim and stem domain of HlgA and HlgB protomers in octameric pore complex. The individual domains (cap, rim, stem, and N-terminal amino latch) and the respective secondary structure components for both the HlgA and HlgB protomers present inside octameric pore complex are indicated in different colours. The distribution and the co-localization of some important amino acid residues in the bicomponent protomeric surfaces for both the HlgA and HlgB protomers are shown in stick model, further represented in different colours.

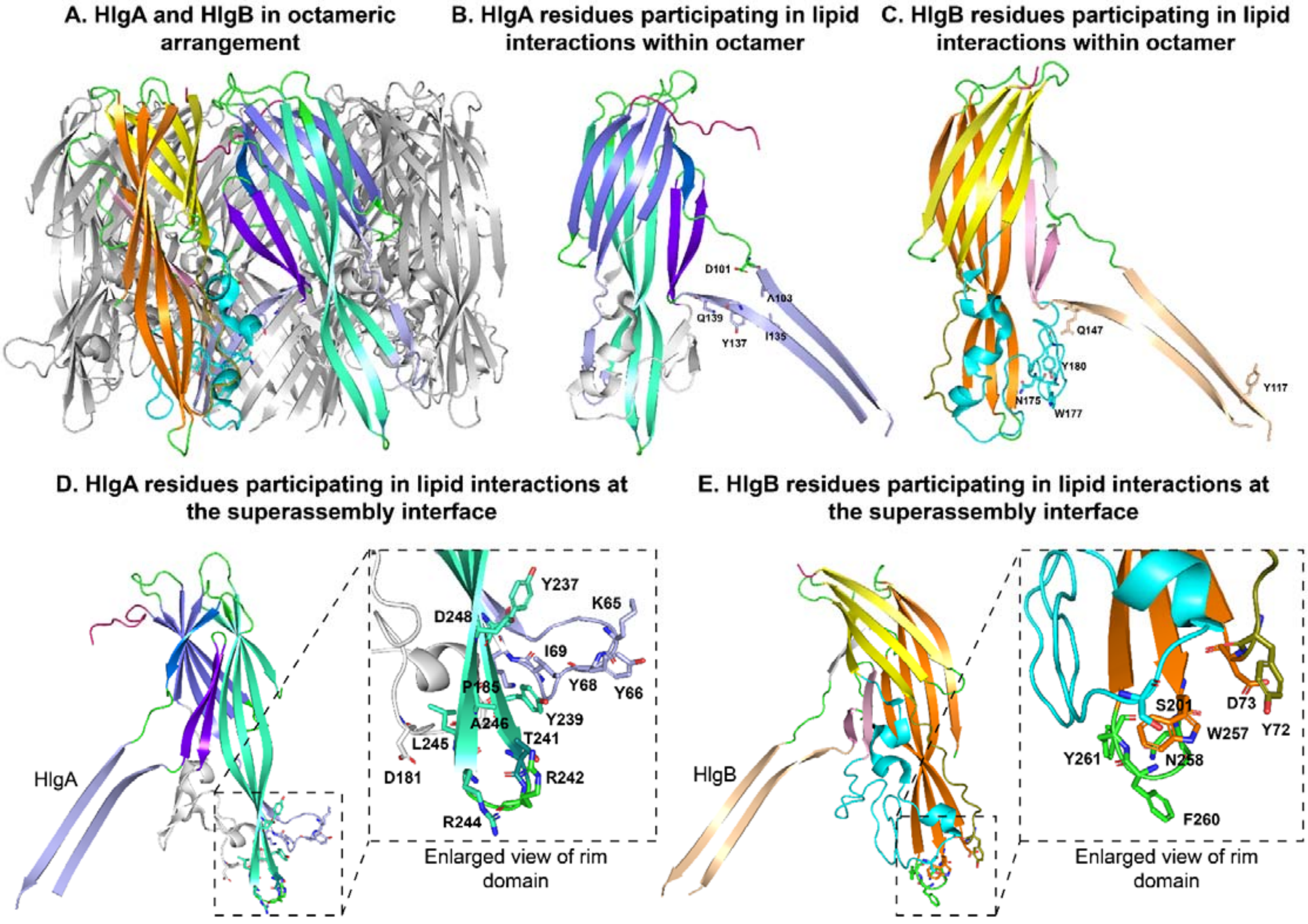


**Figure S17. Identification of crucial amino acids proximal to lipid/cholesterol inside the core of octameric HlgAB pore complex and at superassembly interfaces. A.** The two different protomers of bicomponent hemolysin HlgA and HlgB are shown in two different colours inside the cartoon representation of octameric HlgAB complex (side view). **B.** The residues D101, A103 in β6-hairpin and I135, Y137, Q139 in β7-hairpin, are located solely in stem domain (directed inside cleft of rim and stem domains of HlgA) that are vicinal to lipid/cholesterol in protomeric HlgA. **C.** The amino acids (Y117 in β6-hairpin and Q147 in β7-hairpin) at stem domain and N175, W177, Y180 in large extended disordered segments of rim domain, close to lipid/cholesterol binding pocket in protomeric HlgB are shown (stick model) in wheat and blue respectively. **D.** The residues involved in interfacial interaction in HlgA were located primarily at back-side of MPD/lipid-binding cleft and positioned inside of rim domain, exposed outside part of pore complex. Out of four aromatic sticker residues, Y66, and Y68 belong to small disordered β-turn segments (S55-K70) and Y237, and Y239 located at bottom part of β11 and β12 of rim domain of HlgA respectively. Both hydrophobic (Y66, Y68, and I69 from small, disordered segment, and Y237, Y239, L245, and A246 from β11-β12 turn segment) and polar amino acids (K65 from small, disordered segment turn segment, T241,

R242 and R244 from β11-β12 turn segment, and D181 from large, disordered segment) were shown to co-localize at the close proximal to lipid/cholesterol and responsible for super-assembly interface distribution of protomeric HlgA. **E.** Whereas the crucial amino acids responsible for binding lipid/cholesterol and octahedral HlgAB super-assembly formation were dominated by hydrophobic aromatic amino acid (Y72, W257, F260, Y261) in another protomeric subunit HlgB. In both protomeric subunit, the such co-localization of amino acids predominantly distributed over the bottom part of β11-β12 turn region and small disordered segment (N61-F74) of HlgB. Though the large, disordered segments in both subunits did not possess too many such amino acids, in HlgB, it primarily manifested the MPD/lipid binding pocket.

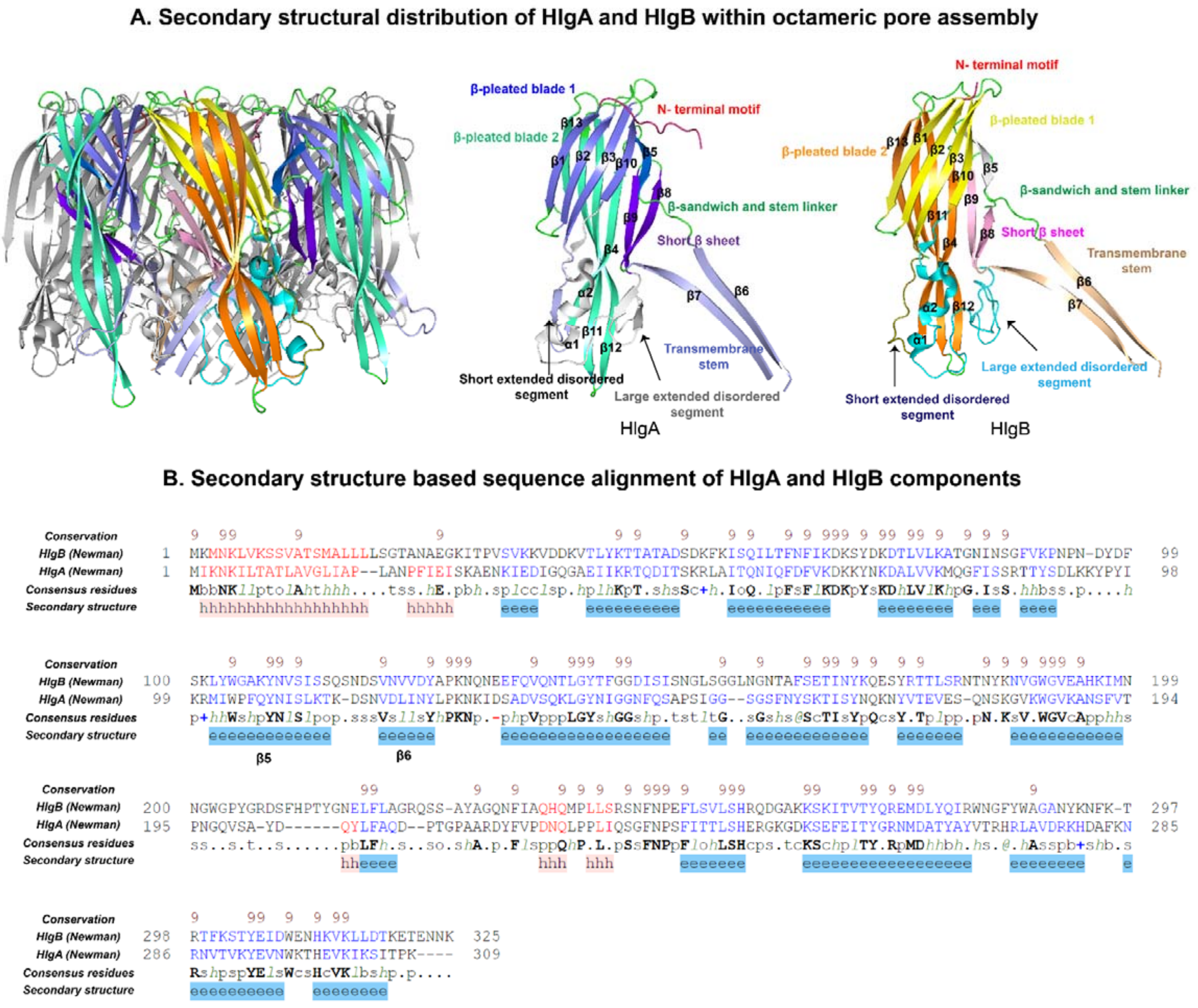


**Figure S18. Molecular architecture of individual bicomponent octameric pore encapsulated inside octahedral super-assembly pore complex. A.** Cartoon representation of γ-hemolysin hetero-octamer (*S. aureus* Newman stain) formed by four components of HlgA and HlgB individually, spanning alternatively through central oligomeric pore. The secondary structural distributions of bicomponent protomers of octameric complex are shown in different colour codes. The overall distribution of structural components of protomeric HlgA (middle) and HlgB (right) that promotes the formation of β-sandwich cap domain, transmembrane β-barrel, and a disordered globule rim domain during oligomerization into octameric pore complex is depicted. Both HlgA and HlgB are composed of a sandwich domain made of β-pleated blade 1 (anti-parallel β-sheet of five short β-strands β1, β2, β3, β10, β5, and a short anti-parallel β-sheet of β9, β8) located on the inner side of cap domain in the pore complex and β-pleated blade 2 (anti-parallel β-sheet of top segments of large extended β-strands β4, β12, β11, & a short strand β13) represents the outer layer of cap domain of pore complex. The transmembrane β-barrel contains four β-hairpins composed of β6, & β7 segments of HlgA and HlgB respectively. The additional structural component including N-terminal amino latch, and β-sandwich cap linker segments are also represented. The disordered globule rim domain of both HlgA and HlgB consists of the bottom part of anti-parallel β-sheet of large extended β-strands β4, β12, and β11; two small helical kinks (α1 & α2); a short, disordered segment from S55-K70 of HlgA and N61-F74 of Hlg; and a large extended unstructured domain from A160-S208 of HlgA, and A169-E225 of HlgB, respectively. **B.** The secondary structure-based multiple sequence alignment of protomeric HlgA and HlgB. The primary sequences of protomeric HlgA and HlgB were aligned with PROMALS3D based on predicted AlphaFold structures for Newman strain. The identical residues of aligned sequences are shown in bold while similar amino acids are shown in red (predicted helix) and blue (predicted β-strand) colour. The adopted secondary structural features are shown with “h” (helix) and “e” (β-strand).

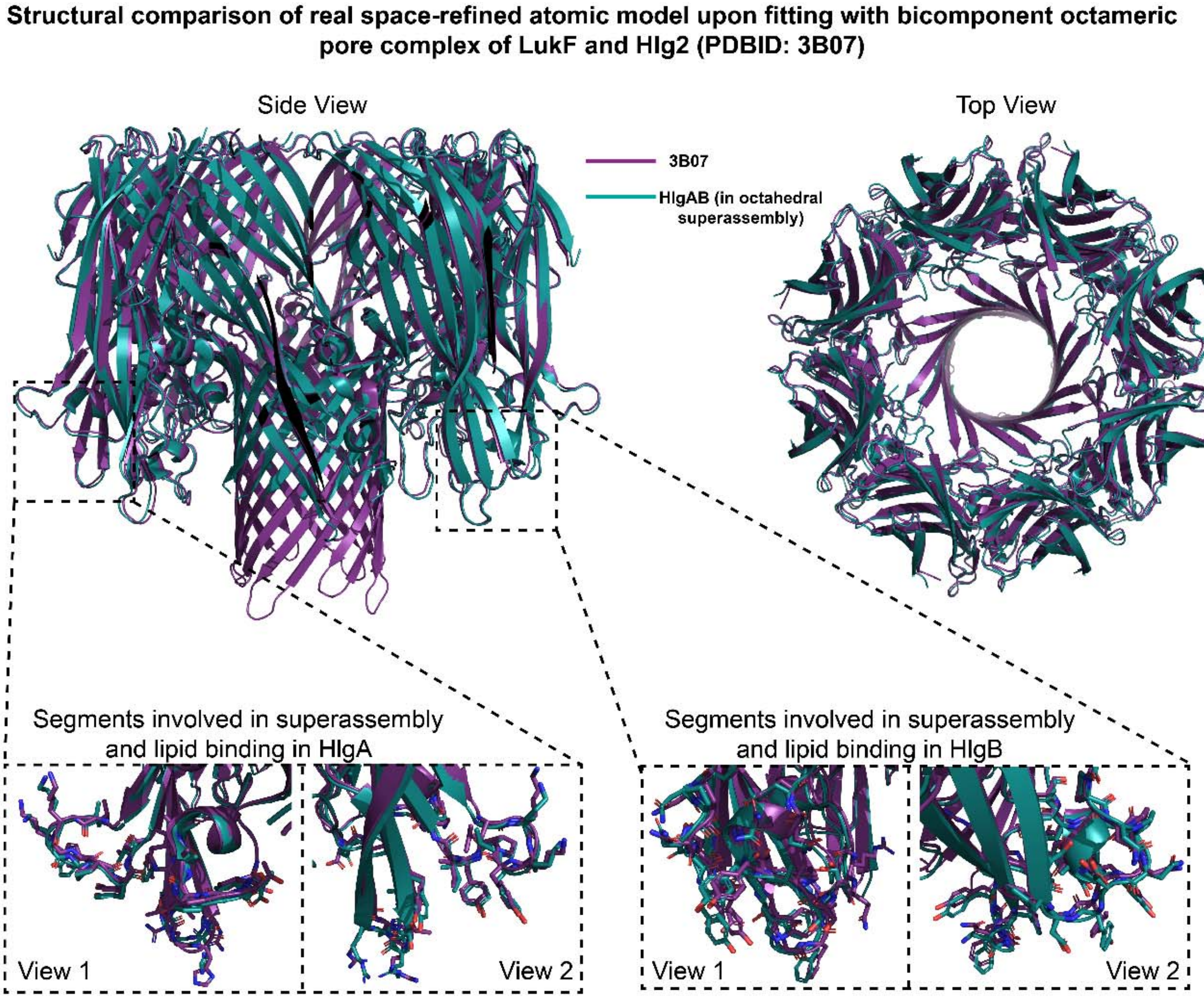


**Figure S19. Structural and conformational comparison of bi-component γ-hemolysin octameric HlgAB pore complex (*S. aureus* Newman strain) present inside octahedral super-assembly pore complex with the molecular architecture of previously reported γ-HL hetero-octamer crystal structure composed of four LukF and Hlg2 respectively (*S. aureus* Mu30 strain).** Side (left) and top views (right) of the cartoon representation of γ-hemolysin octameric HlgAB pore complex (blue) atomic structure (this study), structurally superimposed with γ-HL hetero-octamer of LukF and Hlg2 (PDBID: 3B07). The average RMSD of 15200 cα atoms superimposed with corresponding residues between γ-hemolysin HlgAB octamer and γ-HL hetero-octamer of LukF and Hlg2 are 0.74 Å. The bottom part of transmembrane β-barrel density (β-hairpin segment residues G114-S128 of HlgA, and β-hairpin segment residues G120-N136 of HlgB) was missing for γ-hemolysin HlgAB compared to γ-HL octamer (LukF & Hlg2), while N-terminal segment (K3-A11) of HlgA was clearly resolved via cryo-EM at 3.5 Å resolution. The stereo-representation of residues/segments involved in octahedral super-assembly contacts and the inter-octameric lipid binding for HlgA and HlgB are shown in ball and stick model. The overall structural comparison of the segments responsible for super-assembly contacts and present in the inter-octameric lipid binding interface was 0.532 Å (RMSD) and 0.458 Å (RMSD) for HlgA and HlgB, respectively.

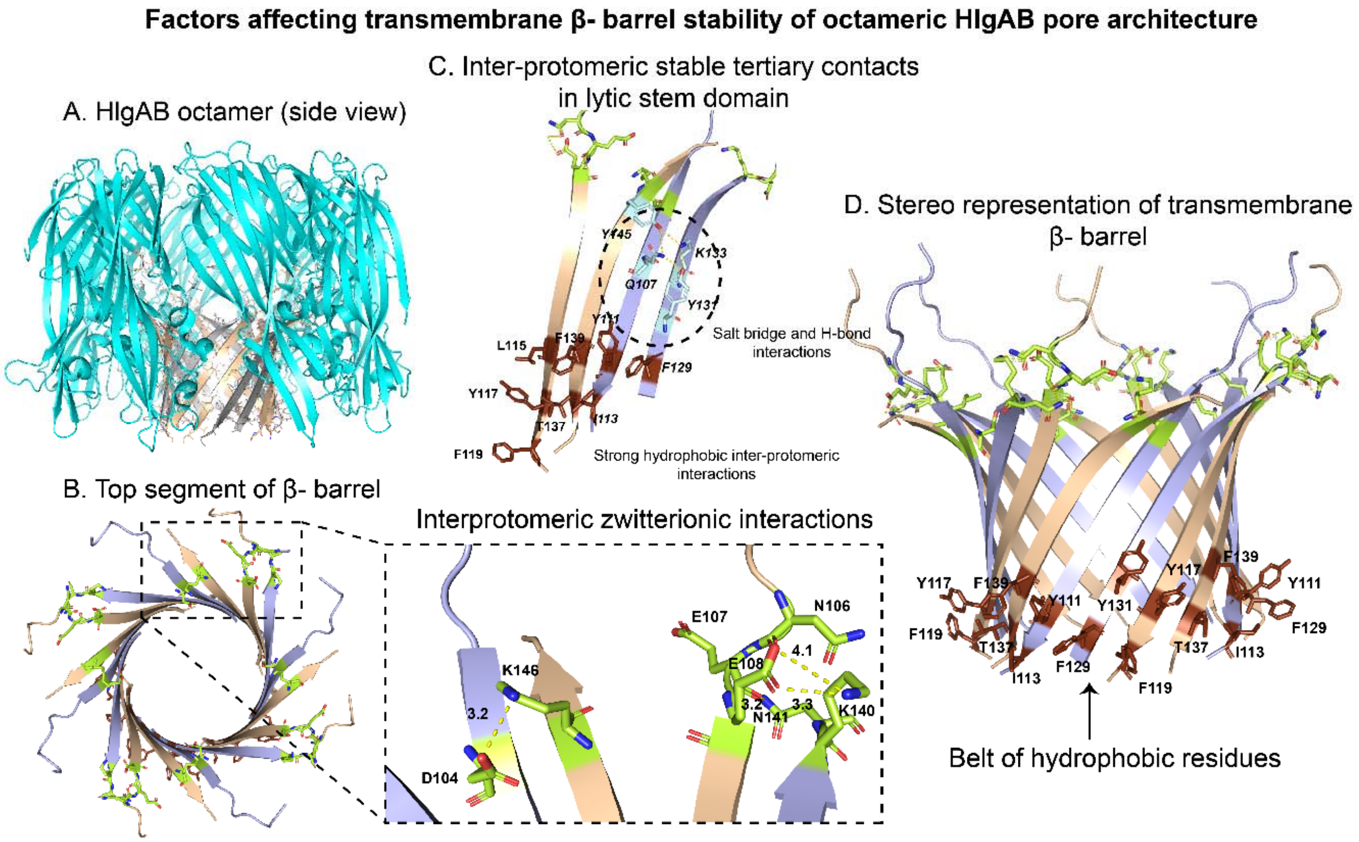


**Figure S20. Transmembrane β-barrel octameric HlgAB pore complex and its conformational stability. A.** Cartoon representation of octameric HlgAB pore complex showing its transmembrane β-barrel domain in silver, composed of four β-hairpins (β6-β7 hairpin) (incomplete hairpin topology) extending from S102-K140 and E107-E148 of HlgA and HlgB respectively. **B.** Stereo-representation of transmembrane β-barrel domain showing terminal segments of β6-strand from β-hairpin of HlgA stabilized by maintenance of ion pairs (lemon green) with nearby terminal segments of β7-strand from β-hairpin of HlgB (D104 of HlgA with K146 of HlgB, and K140 of HlgA with E108 of HlgB), and vice versa. **C.** A hydrophobic clustering (dark chocolate) is involved in intra-protomeric β-hairpin segments (β6 &β7), with key residues being Y111, I113, F129 of HlgA and L115, Y117, F119, T137, F139 of HlgB. Whereas inter β-hairpin clustering of hydrophobic residues were governed by Y111, I113 residues in HlgA, and T137, F139 residues in HlgB. The additional tertiary structural stabilization between the β-hairpin segments of HlgA and HlgB among polar and charge residues Q107, Y131, K133 of HlgA and Y145 of HlgB via hydrogen bond interaction. **D.** A stereo-representation of transmembrane β-barrel domain representing a strong inter β-hairpin hydrophobic belts (dark chocolate) dominated by hydrophobic aromatic amino acids Y111, I113, Y131, F129 of HlgA and Y117, F119, T137, F139 of HlgB residues at the bottom segments of β-barrel whereas the top part gets additional inter-stem ion pair (lemon green) stabilization.

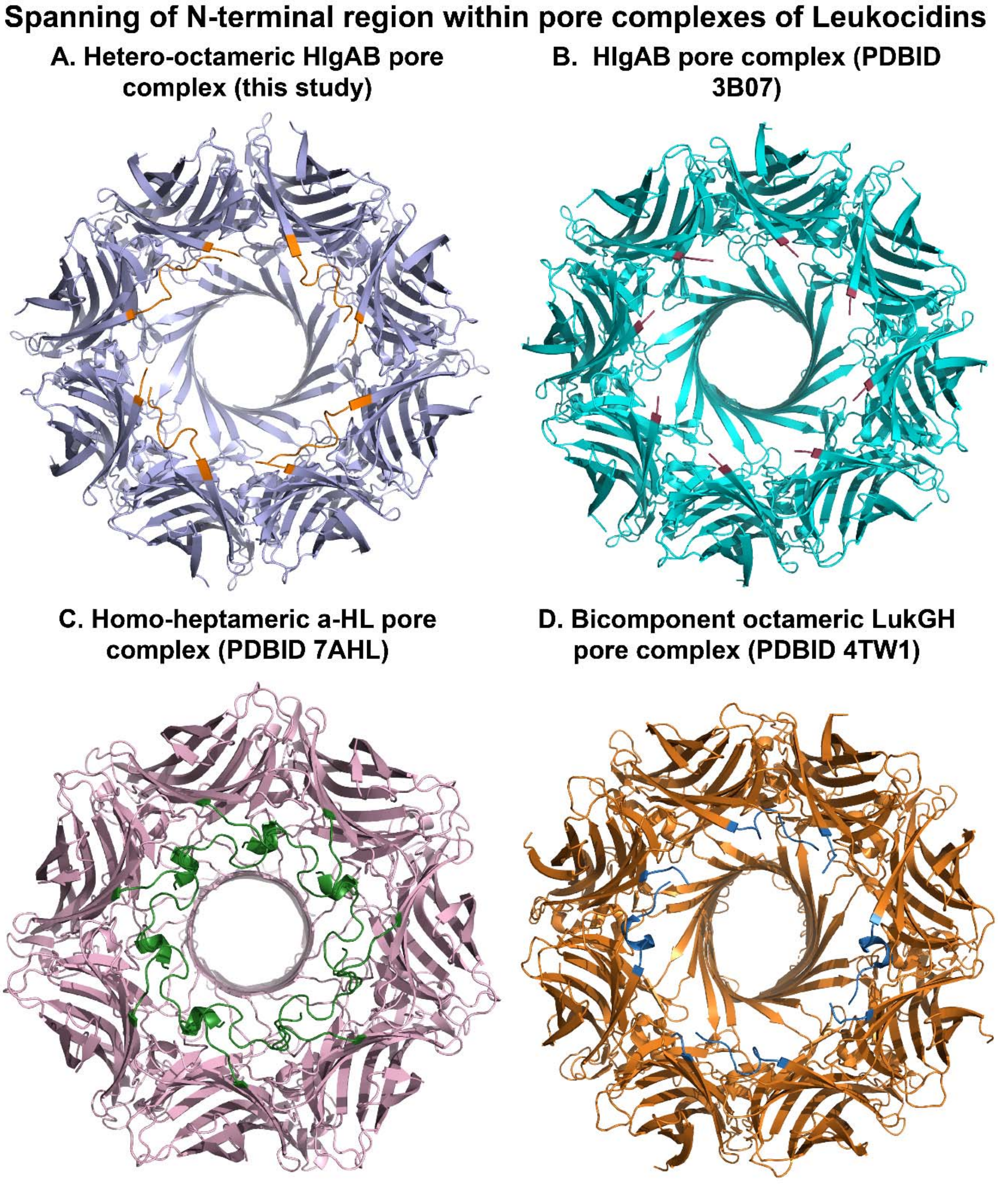


**Figure S21. The representation of N-terminal amino latch segments inside pore complexes formed by four β-PFTs**. **A.** The overall structure, top views of cryo-EM solution state structure of HlgAB octameric pore complex and the N-terminal segments of its constituent individual subunits are represented in orange colour code. **B.** The cartoon-representation of partly resolved N-terminal segments (red chocolate) of octameric MPD-induced crystal structure HlgAB complex (PDBID: 3B07). **C-D.** The spanning of N-terminal segments and the overall structural overviews of two other pore complex; homo-heptameric pore complex of hemolysin (PDBID: 7AHL) and its constituent seven protomeric N-terminal amino latch (green) (C) and bicomponent octameric LukGH (PDBID: 4TW1) transmembrane β-barrel pore complex (D) showing stereo-representation of N-termini (blue) from four adjacent LukG, and LukH protomeric constituents respectively.

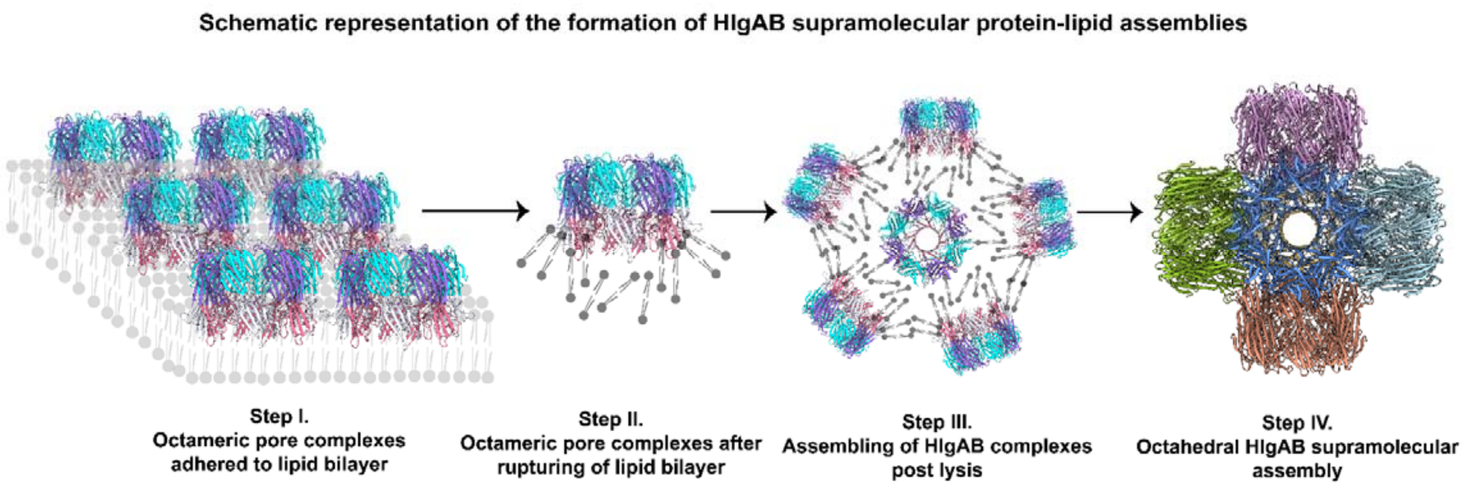


**Figure S22. Schematic representation of the proposed mechanism of formation of HlgAB supramolecular protein-lipid assemblies**. **Step I.** HlgAB toxin components oligomerize and form octameric pore complexes on host lipid bilayer. **Step II.** HlgAB pore complexes rupture the lipid membrane, some lipid moieties remain adhered to the pore complex post membrane lysis. **Step III.** Such pore complexes in proximity associate via their hydrophobic rim domains. **Step IV.** A stable octahedral pore complex of 6 octameric pore complexes provides stability to the rim domains of all 48 toxin components through hydrophobic zipping.

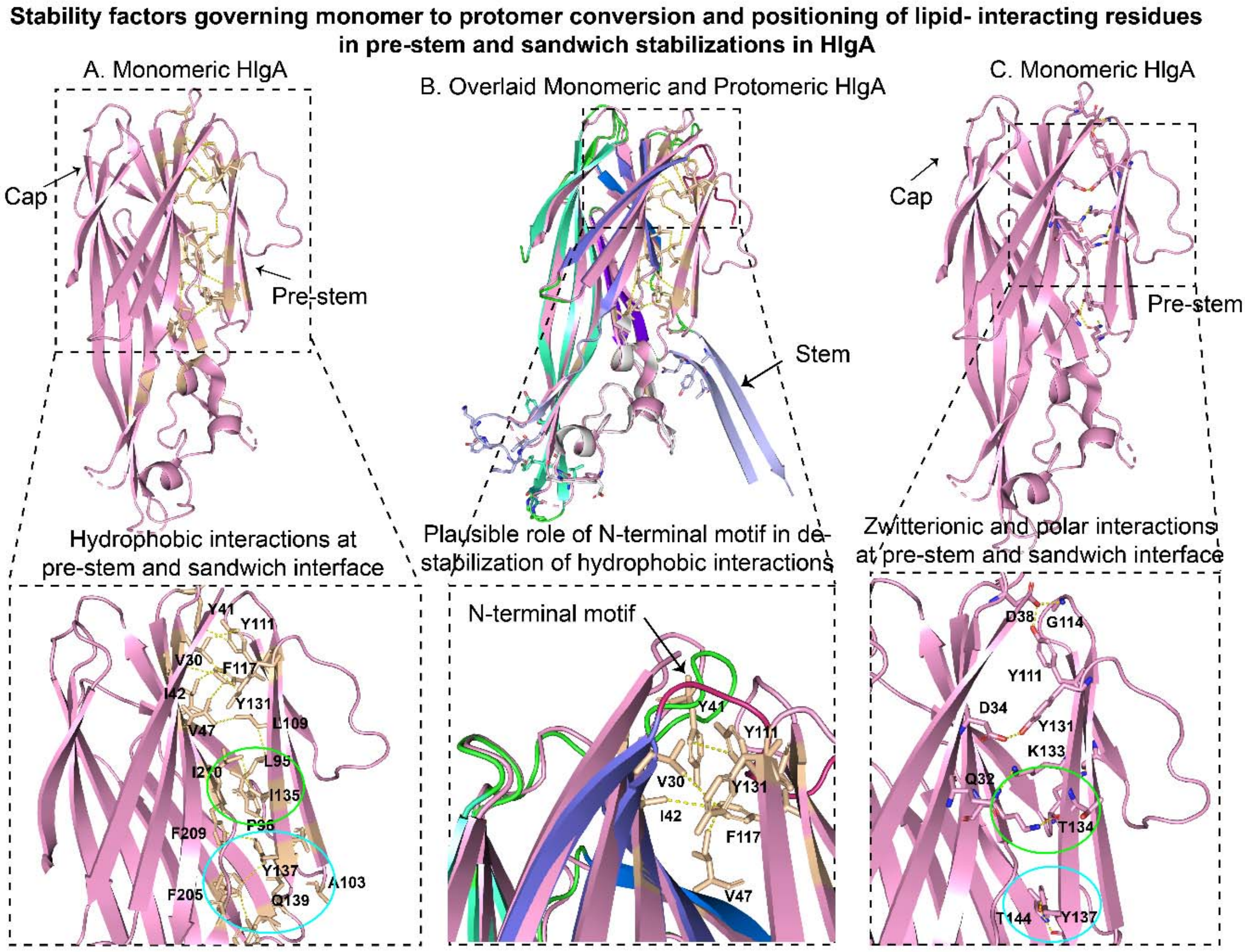


**Figure S23. Structural comparison of protomeric HlgA with its monomeric form and decoding the role of critical lipid/cholesterol binding residues located at stem domain; a plausible role in pre-stem unfolding/displacements from its cap domain. A.** The overall structure of monomeric form of HlgA possessed many hydrophobic amino acids (shown in white) present inside the cap and pre-stem interface. The bottom part of cap and pre-stem interface for monomeric HlgA form a stable hydrophobic cluster (encircled in cyan) of Y137 (located at pre-stem) with F205 and F209 of sandwich cap domain. Whereas the middle part of cap and pre-stem interface form another hydrophobic interaction (encircled in green) of I135 (of pre-stem) with I210 of cap domain. The top part of the cap- pre-stem interface form a large hydrophobic cluster of Y131, Y111, F117 (pre-stem segment) and V30, Y41, I42 (cap domain). **B.** A few polar/charged amino acids has also contributed for ion-pair/dipolar interaction could provide additional stability of cap and pre-stem interface for monomeric HlgA. The inter segment dipolar interactions of Y137 (pre-stem)- T144 (cap domain) at bottom part (encircled in cyan) and T134 (pre-stem)- Q32 (cap domain) at middle part (encircled in green) and Y131 (pre-stem)- D34 (cap domain) and Y111 (pre-stem)- D38 (cap domain) at top part of cap and pre-stem interface for monomeric HlgA is shown. **C.** The protomeric form of HlgA (containing lipid/cholesterol binding residues) was superimposed with its monomeric form and further shown in enlarged view showing the critical positioning of N-terminal amino latch detected from our cryo-EM map causing severe steric effects on nearby hydrophobic and polar residues responsible for monomeric pre-stem and cap interface stabilization. Furthermore, from our study, the amino acid Q139, Y137 and I135 located at stem domain of HlgA protomer in octameric form of HlgAB was found to be close vicinal to lipid/cholesterol density.

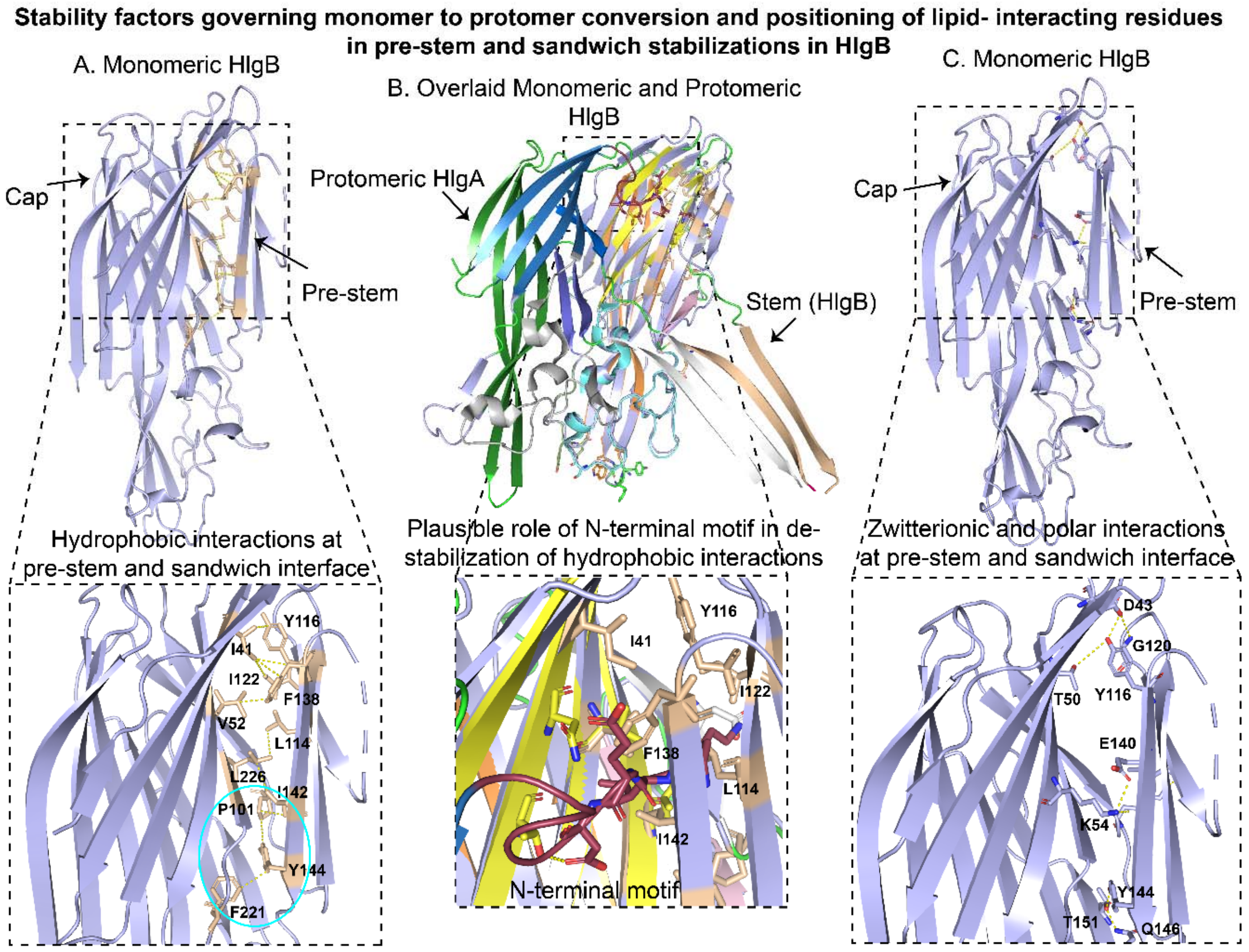


**Figure S24. A detailed structural comparison of protomeric HlgB with its monomeric isoform and decoding the role of critical lipid/cholesterol binding residues located at stem domain; a plausible role in pre-stem unfolding/displacements from its cap domain. A.** The monomeric crystal structure of HlgA contained a layer of hydrophobic amino acids (shown in white) at the cap and pre-stem interface. The bottom segment of cap and pre-stem interface for monomeric HlgB form a hydrophobic interaction between (encircled in cyan) of Y144 (pre-stem) with F221 (cap domain). The middle segment of cap and pre-stem interface get further stabilization by another hydrophobic interaction (encircled in green) of L114 (of pre-stem) with L226 of cap domain. Whereas, in the top segment of the cap- pre-stem interface, many hydrophobic residues of Y116, I122, F138 (pre-stem segment) and I41, V52 (cap domain) are involved to form a large hydrophobic cluster. **B.** A few salt-bridges/dipolar interactions at cap and pre-stem interface for monomeric HlgB were further shown. The inter segment dipolar interactions of Y144 (pre-stem)- T151 (cap domain) at bottom part (encircled in cyan) and

E140 (pre-stem)- K54 (cap domain) at middle part (encircled in green) and Y116 (pre-stem)-D43 and T50 (cap domain) at top part of cap and pre-stem interface for monomeric HlgB is shown in enlarged view. **C.** The protomeric HlgB (containing lipid/cholesterol binding residues) was further superimposed with its monomeric form and shown in enlarged view, showing the critical positioning of N-terminal amino latch from adjacent protomeric HlgA might influenced a strong steric interference on nearby hydrophobic and polar residues of HlgB (monomer) and could cause destabilization at the pre-stem and cap interface. The amino acid Q147 located at stem domain of HlgB protomer was found to be vicinal to lipid/cholesterol density.
