## Supplementary material for "Cryo-EM-based structural insights into supramolecular assemblies of γ-Hemolysin from *Staphylococcus aureus* reveal the pore formation mechanism": Table 1

**Table 1. Cryo-EM data collection, image processing, refinement, and model validation statistics.**

| **I. Data collection and processing** | |
| --- | --- |
| Microscope | Talos Arctica |
| Magnification | x54000 |
| Voltage (kV) | 200 |
| Electron exposure (e-/Å^2^) | 40 |
| Defocus range (μm) | -0.8,-3.0 |
| Pixel size (Å) | 0.92 |
| Number of multi-frame movies | 2287 |
| Number of frames | 20 |
| Imposed symmetry | O |
| Particle number (initial) | 46595 |
| Particle number (final) | 11634 |
| Resolution of electron density map (Å) | 3.55 |
| FSC cut-off | 0.143 |
| Map sharpening B factor (Å^2^) | -129.2 |
| Resolution range (min, 25th percentile, median, 75th percentile, max) | 2.227, 3.137, 3.365, 5.270, 9.086 |
| **II. Model refinement statistics** | |
| Initial model | 3B07 |
| B factor (Å^2^): min/max/mean | 44.19/106.78/66.41 |
| *Model composition* | |
| Non-hydrogen atoms | 104844 |
| Protein residues | 12816 |
| *Root mean square deviations* | |
| Bond length (Å) | 0.007 |
| Bond angle (°) | 0.971 |
| **III. Model Validation** | |
| Molprobity score | 1.77 |
| Clash score | 8.83 |
| CaBLAM outliers (%) | 1.53 |
| Cβ-outliers (%) | 0 |
| Poor rotamers (%) | 0.1 |
| *Ramachandran plot* | |
| Favoured (%) | 95.75 |
| Allowed (%) | 3.91 |
| Outliers (%) | 0.33 |
| Rama-Z score | -0.92 |
| *EMRinger* | |
| EMRinger score | 4.4 |
